# TagF coordinates spike-loading as an intermediate checkpoint in Type VI Secretion System assembly

**DOI:** 10.64898/2025.12.08.692957

**Authors:** Melanie K. Engelin, Pradyot Prakash, Lin Lin, Tobias Mühlethaler, Alexander Heynisch, Joana Pereira, Florian Delbart, Elian M. A. Kuhn, David Baker, Timm Maier, Marek Basler

## Abstract

Diderm bacteria use contact-dependent Type VI secretion systems (T6SS) to gain a competitive advantage within bacterial communities and during infection. Whereas the structural core assembly of T6SS is well characterized, the role of diverse accessory proteins during assembly remains under investigation. One well-conserved accessory protein is TagF, a post-translational inhibitor of T6SS dynamics, which was previously characterized in the context of a kinase-phosphatase signaling relay. Here, we identify a subset of T6SS clusters in which TagF occurs in a distinct regulatory context and without known binding partners. Investigating the role of TagF in the constitutively active *Acinetobacter baylyi* T6SS, we show that though TagF can suppress assembly dynamics, it primarily acts as an assembly coordinator. Using structured illumination microscopy, we show that TagF coordinates the transition from baseplate assembly to sheath elongation by preventing premature and nonproductive sheath assembly in absence of the VgrG spike. Direct interactions with the conserved T6SS tube protein Hcp suggested that TagF blocks sheath elongation by binding and disrupting Hcp hexamers and thus preventing tube formation. Finally, we show that TagF activity depends on crosstalk with TagZ, a previously uncharacterized membrane-associated accessory protein that recruits TagF to the cell periphery, and demonstrate that this interaction can be specifically disrupted by expression of an artificial TagF-binding protein. Together, our findings establish a novel role of TagF as a checkpoint protein that controls for spike-insertion into the baseplate to ensure effective T6SS assembly.

## Introduction

Bacteria often exist in overcrowded niches, where they are in fierce competition over space and resources. Secretion of various specialized proteins has evolved as an effective mechanism to influence their environment. In diderm bacteria, a diverse set of so-called secretion systems export effectors through a variety of mechanisms. The contact-dependent Type VI secretion system (T6SS) is used for predation and nutrient scavenging within bacterial communities, as well as during host-infection. The T6SS has partial homology to other contractile injection systems, and is composed of an envelope-anchoring module, a baseplate, and a contractile sheath-tube assembly (Basler *et al*, 2012; Desfosses *et al*, 2019; Ge *et al*, 2020; Pukatzki *et al*, 2006). T6SS structural components are typically encoded within a single gene locus, along with accessory genes of diverse function (Boyer *et al*, 2009). The T6SS membrane complex, composed of TssM and TssL, anchors T6SS in both membranes, and is often accompanied by additional membrane proteins such as TssJ or TsmK (Durand *et al*, 2015; Felisberto-Rodrigues *et al*, 2011; Kandolo *et al*, 2023; Yin *et al*, 2019). The baseplate is docked onto the membrane complex via twelve TssK-trimers, onto which six TssEF_2_G wedges dock to form the baseplate assembly (Brunet *et al*, 2015; Cherrak *et al*, 2018; Park *et al*, 2018; Zoued *et al*, 2013). The baseplate is the staging module for the insertion of the VgrG_3_/PAAR spike complex, as well as the building platform for sheath-tube assembly (Cherrak *et al*, 2018; Nazarov *et al*, 2018). The spike is decorated with the effector payload: fully folded proteins, often enzymes, that act with high potency even at low doses (Allsopp & Bernal, 2023). The sheath-tube is stacked onto the baseplate in hexameric rings, with the TssBC sheath wrapped around the inner Hcp-tube hexamer (Vettiger *et al*, 2017; Wang *et al*, 2017; Szwedziak & Pilhofer, 2019). Assembly is coordinated by the action of the TssA cap complex, which localizes to the distal end of the sheath-tube during elongation (Santin *et al*, 2018; Schneider *et al*, 2019; Stietz *et al*, 2020; Zoued *et al*, 2017). For translocation of the effector payload, the TssBC subunits rearrange and the entire sheath contracts rapidly, which pushes the Hcp-tube with the spike at its tip through the membrane complex and out of the cell (Szwedziak & Pilhofer, 2019; Vettiger & Basler, 2016; Wang *et al*, 2017). Post-secretion, the AAA+ ATPase ClpV recycles the contracted sheath, and the secreted proteins are resynthesized (Basler & Mekalanos, 2012; Bönemann *et al*, 2009; Kapitein *et al*, 2013; Pietrosiuk *et al*, 2011).

Bacteria use diverse strategies for regulation of timing and localization of their T6SS to ensure successful effector delivery. Some bacteria, such as *Vibrio cholerae*, mainly rely on transcriptional cues to temporally regulate T6SS activity (Manera *et al*, 2021; Proutière *et al*, 2023). Others encode T6SS-accessory genes that coordinate assembly timing and localization post-translationally. The best understood system for fast and effective T6SS deployment is the threonine-phosphorylation pathway (TPP), which was discovered to regulate *Pseudomonas aeruginosa* H1-T6SS. Here, the serine-threonine kinase PpkA, which is activated through the TagQRST membrane damage sensor module, specifically phosphorylates Fha, which drives T6SS assembly in a localized and well-timed manner (Basler *et al*, 2013; Casabona *et al*, 2013; Hsu *et al*, 2009; Li *et al*, 2018; Motley & Lory, 1999; Mougous *et al*, 2007; Silverman *et al*, 2011). To ensure that H1-T6SS assembles exclusively upon PpkA signaling and within a short time frame, the phosphorylated species of Fha is constantly dephosphorylated by the corresponding phosphatase PppA (Silverman *et al*, 2011). Both *Agrobacterium tumefaciens* and *Serratia marcescens* encode homologous TPP systems but differ in both the kinase-activating signal and the downstream phosphorylation target. In *A. tumefaciens* PpkA targets TssL rather than Fha, promoting *Fha-TssL* interactions important for T6SS assembly (Lin *et al*, 2014). In *S. marcescens*, PpkA activation depends on RtkS, but the upstream signal is unclear (Ostrowski *et al*, 2018). In all three characterized T6SS governed by TPP, TagF acts as a conserved post-translational inhibitor of T6SS, and was shown to block assembly by specifically interacting with Fha in both *A. tumefaciens* and *P. aeruginosa* (Silverman *et al*, 2011; Ostrowski *et al*, 2018; Lin *et al*, 2018).

Here, we identified a subset of T6SS clusters that encode TagF proteins in a distinct regulatory context. We show that TagF in *Acinetobacter baylyi* functions in absence of the TPP and plays a key role in the dynamics of the recently discovered T6SS sheath precursors (Lin *et al*, 2022). Furthermore, we show that in *A. baylyi,* TagF functions as a checkpoint protein that stalls T6SS assembly when the VgrG-spike is missing in the baseplate. We determined the structure of TagF and identified putative interaction sites essential for its function. Our biophysical analyses suggested that TagF blocks sheath elongation by directly interfering with Hcp ring integrity and thus tube formation. Moreover, we show that TagZ (ACIAD2698), a previously uncharacterized accessory protein, is essential for TagF recruitment to the membrane to monitor spike assembly. Together, our findings uncover an unexpected role for TagF homologs and provide new insights into a T6SS assembly checkpoint.

## Results

### Genomic context analysis highlights TagF proteins in T6SS clusters without Fha and TPP

TagF was previously shown to be encoded in T6SS clusters together with Fha and the kinase-phosphatase pair PpkA and PppA as part of the TPP, however several clusters were also identified where these accompanying genes were absent (Bayer-Santos *et al*, 2019; Boyer *et al*, 2009; Schwarz *et al*, 2010; Weber *et al*, 2016).

We performed genomic context (GC) analysis on diverse TagF genes using the GCSnap workflow (Pereira, 2021) to identify patterns of co-encoded accessory genes (Supplementary Figure 1A). For downstream analysis, we focused on loci containing at least three of four core T6SS genes (TssM, Hcp, TssC, and ClpV) as indicators for complete T6SS loci. In the resulting dataset of 741 TagF-containing GCs (Figure 1A), almost half of the GCs encoded PpkA, PppA and Fha. These clusters represent the well-characterized TPP previously described in *P. aeruginosa*, *A. tumefaciens*, and *S. marcescens* (Basler *et al*, 2013; Lin *et al*, 2018; Ostrowski *et al*, 2018; Silverman *et al*, 2011). Surprisingly, 22.0% of all analyzed GCs encoded neither the phosphatase-kinase pair nor Fha (163 GCs). Using GCSnap, we categorized the data by hierarchical clustering based on gene content of the respective GCs. Because structural genes were consistently present across T6SS loci in our dataset, the clustering primarily reflected variation in associated accessory gene content. This analysis revealed a distinct cluster comprising 47 functional GCs exclusively coding for *tagF* but lacking *fha* or TPP components (Supplementary Figure 1B). In this operon type, a specific subset of T6SS accessory genes, referred to as T6SS Accessory Gene Set A (TSA), was enriched with homologs for *tagN* (100.0%), *tagX* (100.0%), *tslA* (83.0%), *tsmK* (42.6%), and an uncharacterized gene (85.1%; later named *tagZ*), all of which were previously described in *Acinetobacter* (Kandolo *et al*, 2023; Lin *et al*, 2019a, 2022; Ringel *et al*, 2017; Weber *et al*, 2016).

**Figure 1:**
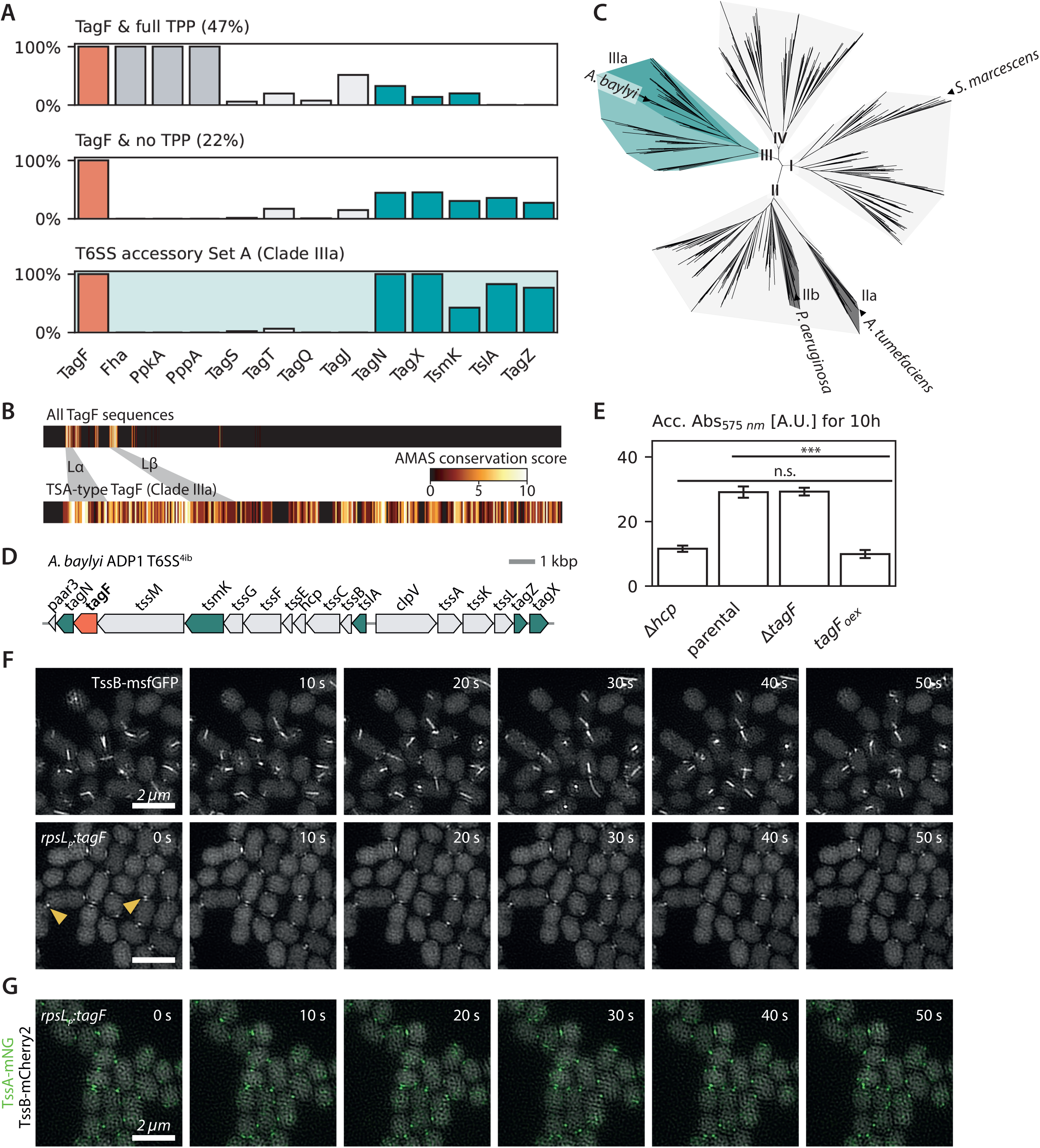
*A. baylyi* TagF is part of the novel conserved regulatory context TSA and a functional inhibitor of T6SS dynamics. **A** Bioinformatic analyses on TagF GC, shown is a comparison of relative frequency of accessory gene occurrence in GCs with intact TPP and without. The subset of GCs encoding TSA is shown in the bottom panel. **B** MSA of all TagF sequences highlights overall sequence conservation of α and β loops at the N-terminus. TagF sequences co-encoded with TSA show overall high sequence conservation (Supplementary File 3). **C** Phylogenetic analysis of TagF sequences from GC analysis visualized using iTOL. Sequences cluster in four clades, clades I, II, IV contain TagFs co-encoded with variable sets of TPP genes. Clade III is enriched for TagF proteins without TPP context, and clade IIIa corresponds to TagFs encoded in a TSA-context. **D** *A. baylyi* ADP1 T6SS cluster; *tagF* in orange, TSA-type accessory genes in green, structural genes in grey. **E** CPRG *E. coli* lysis accumulation assay comparing T6SS activity of *A. baylyi* strains with a *tagF* deletion mutation and a strain overexpressing TagF (*TagF_oex_)* to a T6SS-negative *Δhcp* mutant and the parental strain. **F** Time-lapse 2D-SIM following sheath dynamics (TssB-sfGFP) of the parental strain and TagF_oex_. Arrows highlight examples of stalled sheath precursors at contact sites in *TagF_oex_*. **G** 2-color time-lapse 2D-SIM imaging *TagF_oex_* sheath (TssB-mCherry2) and TssA-mNG localization. Single channels in Supplementary Figure 4.

Together, our analyses show that *tagF*-containing T6SS clusters encode variable sets of accessory genes. We define TSA, a novel T6SS regulatory context that exclusively contains *tagF* homologs lacking *fha*/TPP but instead encode a distinct set of accessory genes that remain largely uncharacterized.

### TagF homologs have conserved N-terminal and diverse C-terminal domains

Building on the distinct accessory gene composition between TSA and TTP, we hypothesized that changes in potential interaction partners might drive divergence in TagF protein sequences as a structural or functional adaptation.

Multiple sequence alignments (MSA) of 655 TagF protein sequences using MUSCLE (Edgar, 2004) revealed strong conservation in the N-terminal region. Specifically, two conserved hotspots, corresponding to the α and β loops in the crystal structure of *P. aeruginosa* PaTagF1, were highly conserved across all homologs (Figure 1B) (Lin *et al*, 2018). These loops include residues previously shown to be essential for TagF function and Fha binding in both *P. aeruginosa* H1-T6SS and *A. tumefaciens* (Lin *et al*, 2018). Importantly, these residues remained conserved even in TSA-type TagF homologs.

To explore how regions beyond the conserved N-terminal loops show sequence divergence, we built a maximum-likelihood phylogenetic tree from the MSA and annotated each leaf with the accessory gene content of its T6SS locus, linking protein variation to regulatory context (Kumar *et al*, 2024; Letunic & Bork, 2024)(Figure 1C, Supplementary Figure 2). Overall, we identified four major clades, in which the clustering of leaves partially reflected the genomic context of the corresponding TagF homologs. Previously characterized TagF homologs were present in clades I and II, with *Serratia* TagF homologs in clade I, the *A. tumefaciens* (*A. fabrum*) TagF-PppA fusion in subclade IIa and *P. aeruginosa* TagF subclades IIb. TagF variants in clade III, including those from *Acinetobacter* species, lacked both Fha and TPP. Importantly, homologs in subclade IIIa recapitulated the accessory gene set characteristic of TSA locus type identified in prior GCSnap results. A focused MSA of this group revealed that the C-terminal region remains highly conserved (Figure 1B).

In summary, all TagF homologs share a conserved N-terminal region but diverge in their C-termini. During phylogenetic analysis, we identified subclade IIIa, which contained TagF homologs encoded in TSA-specific contexts. Taken together, our findings suggest that these TagF homologs may represent a novel class with a shared mechanism of action.

### Overexpression of TagF arrests T6SS assembly in *Acinetobacter baylyi*

To investigate TagF function within the novel regulatory context of TSA, we chose *Acinetobacter baylyi* ADP1 as a model to due to its natural competence and well-characterized T6SS dynamics (Figure 1D)(Ringel *et al*, 2017; Lin *et al*, 2019b, 2019a, 2022).

Initially, we tested the effect of *A. baylyi* TagF on T6SS activity by monitoring lysis of susceptible LacZ+ *E. coli* (Ringel *et al*, 2017). Cumulative release of LacZ from lysed prey cells and the subsequent conversion of the chromogenic substrate over the first 10 h of competition was used as a measure for T6SS activity. This allowed us to obtain a high dynamic range and sensitivity to delayed killing dynamics observed in some mutants. Using this assay, we determined that T6SS dependent lysis of *ΔtagF* was comparable to the parental strain (Figure 1E, Supplementary Figure 3), as shown previously (Ringel *et al*, 2017).

Next, we used the strong constitutive promoter of *rpsL* to overproduce TagF (labeled as TagF_oex_) from a neutral site on the genome, to test if it can block T6SS assembly at higher expression levels. Whole proteome quantitative mass spectrometry showed that TagF expression was increased about 70-fold in TagF_oex_, while other T6SS proteins, except Hcp, exhibited comparable levels of expression relative to the parental strain (Supplementary File 4). Surprisingly, TagF overexpression completely abrogated T6SS activity, with no significant increase in prey lysis of TagF_oex_ above the T6SS-defective *Δhcp* mutant (*p* = 0.15).

To assess the effect of TagF_oex_ on T6SS dynamics, we additionally performed two-dimensional structured illumination microscopy (2D-SIM) to localize sheath (TssB-sfGFP) in live cells (Figure 1F, Supplementary Video 1). No sheath dynamics were observed for TagF_oex_ in 3 min long time lapses (3 biol. replicates, 6 fields of view (FOV)). Interestingly, diffraction limited sheath foci accumulated in TagF_oex_ at the cell periphery at the cell-cell contacts over time. Previously, similarly arrested contact-dependent sheath foci were shown in *A. baylyi* lacking *tagX* (Lin *et al*, 2022). To determine the composition of the arrested T6SS structures, we also analyzed 2D-SIM imaging of TssA-mNG localization in TagF_oex_ and found that TssA-mNG foci localized to cell-cell contacts as well (Figure 1G, Supplementary Figure 4).

Taken together, these results indicate that the TagF homolog in *A. baylyi* is a functional post-translational inhibitor of T6SS dynamics. Importantly, the observed sheath and TssA localization to cell-cell contacts suggests that overexpression of TagF arrests T6SS assembly at a precursor state prior to sheath elongation, which already contains essential components required for the full assembly.

### TagF controls checkpoint that prevents sheath assembly on incomplete baseplate

To define the target of TagF regulation at native expression levels, and building on the observation that TagF_oex_ produces T6SS assembly precursors, we systematically imaged baseplate deletion mutants using 2D-SIM to identify those, where assembly would stop at a state similar to TagF_oex_ (Supplementary Figure 5A). In agreement with previous reports, no sheath foci were detected in any mutants of *tssK* or the *tssF/G* baseplate wedge genes (Lin *et al*, 2022). We observed sheath foci in *ΔtssE*, but they were not stable over time.

Surprisingly, we detected stable sheath foci in *ΔvgrG1-4* and *Δhcp* (Supplementary Figure 5B). We then deleted *tagF* in these strains to test the role of TagF in forming sheath foci. No change was observed in *ΔhcpΔtag*F compared to parental Δ*hcp* strain (Supplementary Figure 5C). However, deletion of Δ*tagF* in Δ*vgrG1-4* background resulted in formation of short sheath structures with clear assembly, contraction, and recycling dynamics (Figure 2A-C, Supplementary Figure 6, Supplementary Video 2). T6SS assembly frequency of *ΔvgrG1-4ΔtagF* (0.204 ± 0.03 assemblies/cell*min; *n* = 240 cells) was reduced to 31% of the activity of the strain expressing VgrGs and TagF (0.653 ± 0.106 assemblies/cell*min; n = 284 cells). Importantly, deletion of *hcp* in *ΔvgrG1-4ΔtagF* blocked assembly of elongated sheaths (Figure 2D-F, Supplementary Figure 7), suggesting that the sheath assembly in *ΔvgrG1-4ΔtagF* is dependent on Hcp.

**Figure 2:**
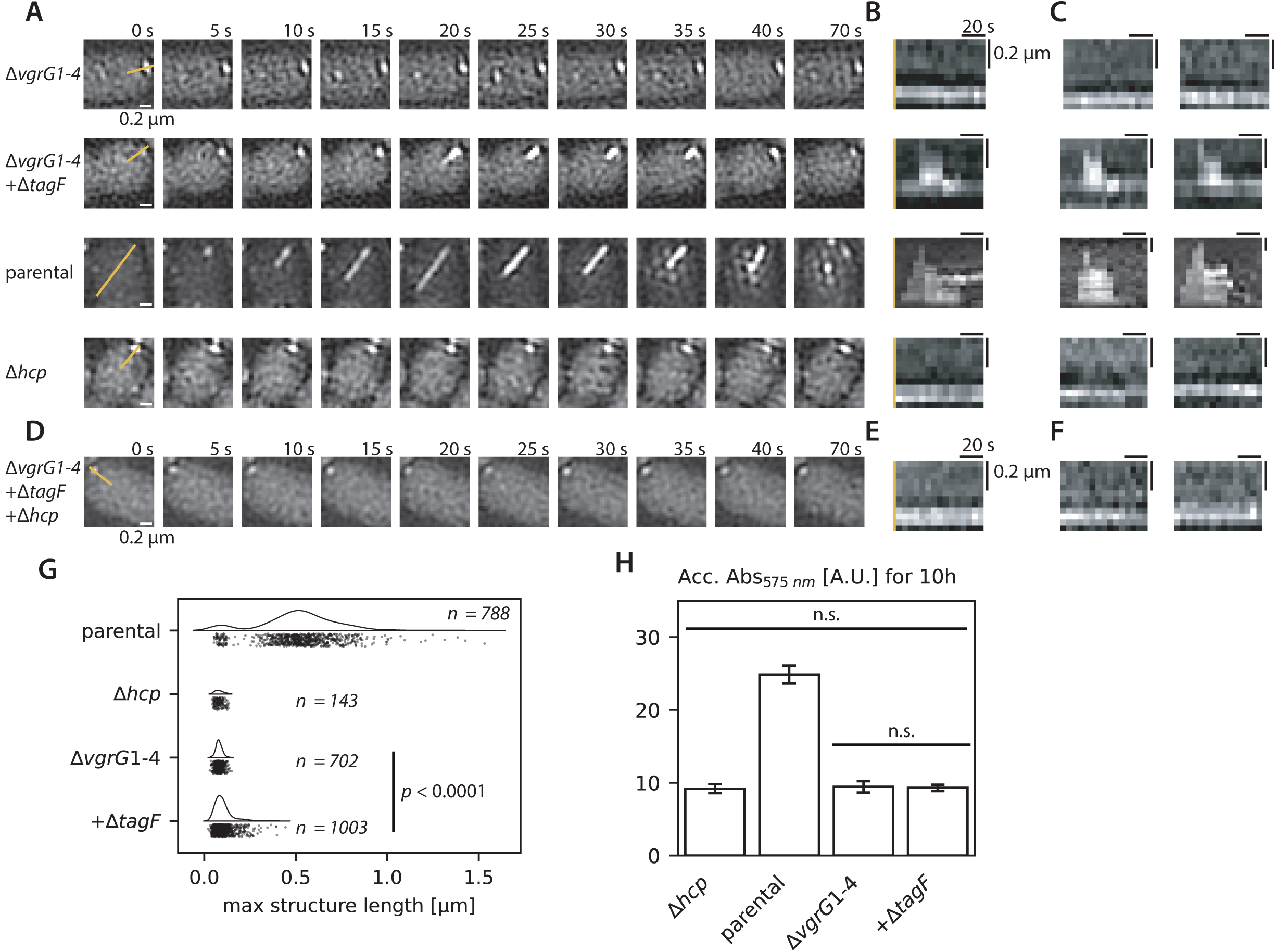
T6SS precursors as a regulatory checkpoint. **A** Representative 2D-SIM time-lapse following sheath dynamics (TssB-sfGFP) of single *A. baylyi* cells from different mutant strains highlighting TagF function. Top to bottom: *ΔvgrG1-4*, *ΔvgrG1-4ΔtagF* mutants in comparison to parental strain and Δ*hcp*. Yellow lines in the first panels indicate kymograph origin in (B). **B** Kymographs of the time-lapse imaging in (A), highlighting the differences in sheath dynamics between strains. **C** Additional representative kymographs, further examples in Supplementary Figure 6 and Supplementary Figure 7. Scale as in (B). **D** Representative 2D-SIM time-lapse showing loss of sheath dynamics (TssB-sfGFP) in *ΔvgrG1-4ΔtagFΔhcp*. **E** Kymographs of the time-lapse imaging in (D). **F** Additional representative kymographs, scale as in (B) **G** Violin plots showing image analysis of maximum sheath structure lengths observed between the strains in (A). Shown are cumulatively the analysis from 3 biological replicates with each 3 FOV. Statistical analysis: Kruskal-Wallis-ANOVA; Dunns’ multiple comparisons test. **H** CPRG *E. coli* lysis accumulation assay comparing T6SS activity of *A. baylyi* strains shown in (A). Tested for statistical significance using ANOVA.

We next measured the maximum length of sheath structures for the observed assemblies in the different mutant backgrounds (Figure 2G). Of the analyzed sheath structures in *ΔvgrG1-4ΔtagF* (n = 1003), 11.5% extended beyond 150 nm (n = 103) and only 5.6% extended past 200 nm (n = 56). Overall size distribution of the dynamic assemblies in *ΔvgrG1-4ΔtagF* was significantly different from the static sheath foci in Δ*vgrG1-4* (*p* < 0.0001, Dunns’ multiple comparisons test; Kruskal-Wallis-ANOVA) and Δ*hcp* (*p* = 0.0007), which remained diffraction limited. In contrast, in the parental strain, out of 788 observed assemblies, half of the assembled sheath structures reached at least 519 nm (n = 394). Moreover, the short dynamic sheath structures failed to kill prey cells as neither *ΔvgrG1-4*, nor *ΔvgrG1-4ΔtagF* showed any *E. coli* lysis above the T6SS negative control *Δhc*p (Figure 2H).

Taken together, these results suggest that TagF arrests T6SS assembly, and sheath elongation can only proceed once the VgrG-spike is located within the baseplate. Deletion of *tagF* in a background lacking all VgrG proteins bypassed this checkpoint and resulted in short dynamic, but unproductive, sheath assemblies.

### TagF structure-function analysis reveals interaction sites

We purified and crystallized *A. baylyi* TagF to gain further insight into both structural and mechanistic details of TagF function. TagF crystallized at 1.8 Å resolution in space group P2_1_2_1_2_1_. The asymmetric unit coordinated a TagF monomer with 11 cadmium ions from the crystallization condition, which were used to phase the structure experimentally. The structure revealed a two-domain architecture, featuring N- and C-terminal domains with distinct α+β and α/β folds (Figure 3A, Supplementary Figure 8). Extensive inter-domain interactions stabilize the compact fold of the protein.

**Figure 3:**
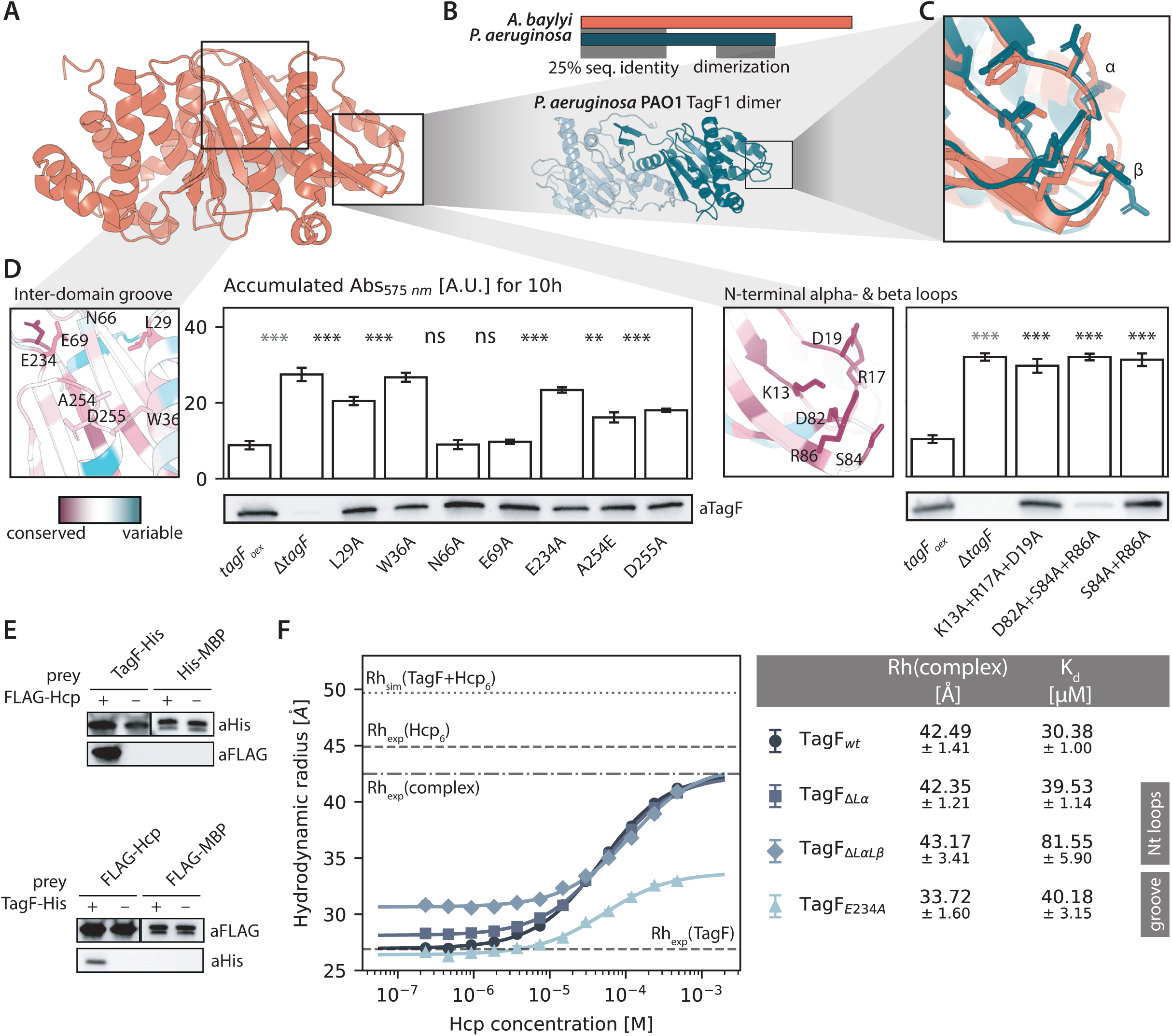
*A. baylyi* TagF structure and interaction screens. **A** Side view of *A. baylyi* TagF X-ray structure (PDB: 9T0N). **B** Summary of pairwise sequence alignment analysis for *A. baylyi* and *P. aeruginosa* TagF (Supplementary File 3). Below: *P. aeruginosa* PaTagF1 dimeric X-ray structure shown for comparison (PDB 2qnu). Monomers shown in different shades of blue. **C** Detailed view of α and β loop structural alignment between both structures, RMSD = 1.41 over 32 residues. **D** Sequence-structure conservation analysis using ConSurf highlights two distinct putative interaction sites. Residues of interest tested using the mutagenesis scanning approach are shown in stick representation. *E. coli* lysis accumulation assay comparing T6SS activity of *A. baylyi* TagF_oex_ point-mutant strains. Below: Western blot analysis of TagF mutant expression from the *A. baylyi* soluble lysis fraction using specific anti-TagF polyclonal antibody. **E** Affinity-tag based co-purification from *E. coli* using TagF-His and FLAG-Hcp identifies a direct and reciprocal interaction. **F** FIDA analysis of TagF-Hcp interaction by measuring change in Rh to follow complex formation. Shown are TagF as well as loop and groove point mutants. Experimental Rh for pure TagF and Hcp_6_ plus simulated Rh for TagF+Hcp_6_ are indicated with dashed lines. Rh(complex) for TagF wildtype is indicated with dot-dashed line. Panel to the right details Rh(complex) and K_d_ for each sample with standard deviation.

In agreement with our sequence alignments, the N-terminal domain TagF_1-150_ showed a high degree of structural homology to the *P. aeruginosa* PaTagF1 crystal structure (Filippova *et al*, 2007) (Figure 3B). Remarkably, the two surface exposed loops α and β of *A. baylyi* TagF showed strong structural conservation to PaTagF1, corresponding to the putative Fha1 interaction site in *P. aeruginosa* (Lin *et al*, 2018) (Figure 3C). In contrast, the C-terminal TagF_151-320_ domain featured an α/β fold with no homology to PaTagF1 or other characterized proteins. We then visualized sequence conservation between all identified TSA-specific TagF homologs on the *A. baylyi* structure to gain further insight into distinct conserved features and potential interaction hotspots (Supplementary Figure 9AB). Next to the α- and β-loops universally conserved in all known TagF homologs, we identified an additional surface exposed region of high sequence conservation on the protein surface: A groove lined with negatively charged residues located at the interface between N- and C-terminal domain, which is absent on the PaTagF1 structure.

To test if the conserved residues are important for TagF function, we replaced them with alanine and overexpressed the mutated proteins in *A. baylyi* (Figure 3D, Supplementary Figure 9C). We found that mutations in the α- (K13A/R17A/D18A) and β- (R84A/R86A) loops directly impacted TagF function, resulting in a loss of TagF-mediated inhibition of T6SS assembly without affecting TagF protein stability. In the groove, we identified two hydrophobic residues in helix 1 of the N-terminal domain, L29 and W36, likely coordinating positioning of helix 1, and two charged residues in the C-terminal domain at the opposite side of the groove, E234 and D255, where point mutations in TagF abolished its ability to block T6SS upon overexpression. Interestingly, introduction of a negative charge in the center of the groove via a A254E point mutation also resulted in loss of TagF-mediated inhibition on T6SS activity, while residues E69 and N66 played no role.

In summary, we discovered a high degree of structural conservation in the N-terminal domain between PaTagF1 and *A. baylyi* TagF. Surprisingly, residues in the α and β loops are essential for TagF function in *A. baylyi*, which lacks Fha, suggesting a different interaction partner. Conserved residues in the groove on *A. baylyi* TagF present a putative interaction site specific to this novel class of TagF proteins.

### Direct TagF-Hcp interaction destabilizes Hcp ring structure

Having mapped potential TagF interaction interfaces, we performed co-purifications to find potential TagF interaction partners. *A. baylyi* cell lysate solubilized in 1% *n*-dodecyl-beta-D-maltoside (DDM) and clarified by ultracentrifugation was incubated for 60 min at 4°C with purified TagF protein trapped on magnetic sepharose beads. Co-purified proteins were identified using quantitative LC-MS/MS. In three independent experiments, we consistently identified Hcp as significantly enriched in the sample with TagF bait (43.6-fold over 3 biol. reps, adj. *p* = 1.29E-06, Supplementary Figure 10A). This interaction was further confirmed by performing reciprocal small-scale purifications of TagF and Hcp in expressed *E. coli* (Figure 3E, Supplementary Figure 10BC). Importantly, we only co-purified FLAG-Hcp with immobilized TagF-His, and TagF-His with immobilized FLAG-Hcp, indicating that TagF and Hcp can interact specifically and directly, and in absence of additional factors.

To further characterize the interaction between TagF and Hcp, we isolated both proteins from *E. coli* and measured the binding affinity using flow-induced dispersion analysis (FIDA) (Supplementary Figure 11AB). The experimental hydrodynamic radius of TagF was determined as Rh(TagF) = 29.3 ± 0.20 Å consistent with a monomeric protein. The hydrodynamic radius for Hcp Rh(Hcp) = 44.5 ± 0.36 Å suggested a hexameric assembly. This agreed with both mass photometry of Hcp indicating hexameric complex, and negative stain electron microscopy (EM) revealing typical ring-shape topology similar to other Hcp_6_ (Supplementary Figure 11CD) (Ruiz *et al*, 2015; Wang *et al*, 2017).

To determine the TagF-Hcp interaction parameters, FIDA was used to track the hydrodynamic radius of fluorescently labelled TagF in presence of increasing concentrations of Hcp_6_ (Figure 3F). Interestingly, the hydrodynamic radius of the TagF-Hcp Rh(complex) = 42.5 ± 1.4 Å was already smaller than a free Hcp_6_ ring, and much smaller than the simulated Rh for the TagF-Hcp_6_ complex (Rh_sim_ = 49.7 Å). Together, this suggested that TagF does not bind intact Hcp_6_ but instead breaks the ring-structure for complex formation, however, the exact composition of the complex is unclear. Importantly, the dissociation constant of the TagF-Hcp interaction could be determined at K_d_ = 30.4 ± 1.0 µM. Based on the precise cellular quantification of T6SS proteins as reported in Lin *et al*, 2019, Hcp concentration exceeds TagF 23-fold. This indicates that up to 90% of TagF would be associated with Hcp, while Hcp_6_ remains in excess and available for sheath-tube assembly.

We next sought to identify which TagF interaction site, the N-terminal α+β loops or the groove, would drive Hcp binding. We purified key mutants previously tested in *A. baylyi* (Figure 3D) and repeated the FIDA assay to assess Hcp_6_ binding (Figure 3F).

Surprisingly, only a double-loop mutant TagF_ΔLαLβ_, carrying a total of five point mutations, significantly impacted affinity. Unexpectedly, we found that a mutant in the groove region produced a smaller TagF-Hcp complex, while preserving affinity.

In summary, we present several lines of evidence that *A. baylyi* TagF directly interacts with Hcp. FIDA suggests a moderate affinity for complex formation, with a strong indication that TagF can disrupt the Hcp_6_-ring upon binding, both Lα and Lβ as well as the groove contribute, but neither binding site dominates Hcp binding.

### TagZ is required for TagF mediated T6SS regulation

Based on the affinity of the TagF-Hcp interaction measured *in vitro*, additional mechanisms are likely needed to locally increase TagF concentration at T6SS precursors within cells to fully block assembly in the absence of the spike. Therefore, we investigated if additional TSA-associated genes would contribute to TagF function. We overexpressed TagF in single deletion strains of all other five accessory genes within the cluster and screened for increase in *E. coli* lysis compared to TagF overexpression in the intact parental strain. Interestingly, compared to TagF_oex_, we only observed a significant increase in T6SS activity for a *ΔtagZ* (*ΔACIAD2698*) mutant overexpressing TagF (*p* = 0.00018), whereas deletion of *ΔtagX*, *ΔtsmK*, *ΔtagN*, *ΔtslA* had no impact on the TagF_oex_ phenotype (Figure 4A, Supplementary Figure 12ABC, Supplementary Video 3).

**Figure 4:**
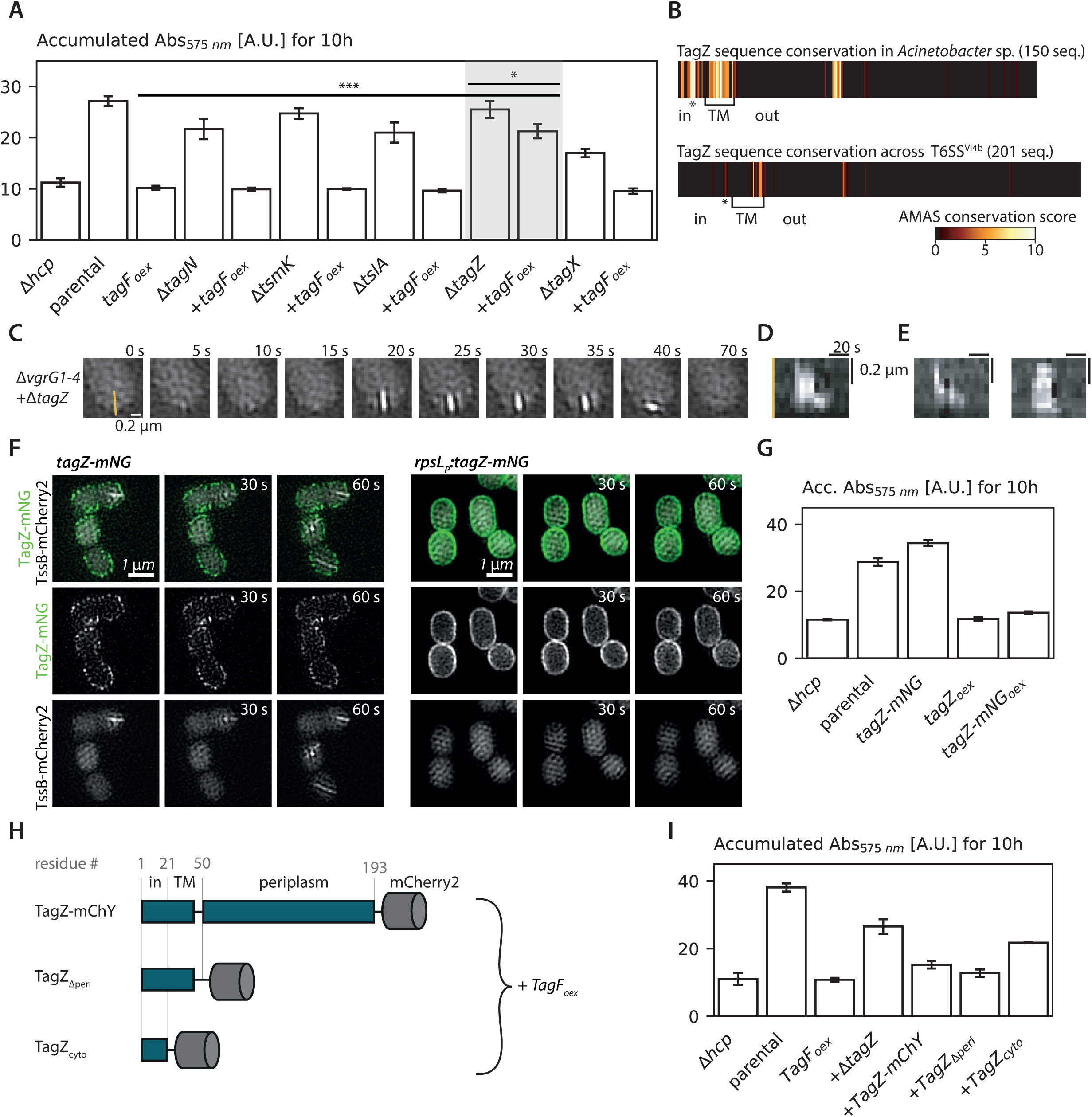
TagZ periplasmic domain is non-essential for complementation of the TagF phenotype. **A** CPRG accumulation assay comparing T6SS-dependent *E. coli* lysis of strains overexpressing TagF in accessory gene deletion backgrounds. **B** MSA of all TagZ sequences from *Acinetobacter* and all identified TagZ from TSA-clusters. Predicted TagZ topology as indicated. **C** Representative 2D-SIM time-lapse showing loss of sheath dynamics (TssB-sfGFP) in *ΔvgrG1-4ΔtagZ*. **D** Kymographs of the time-lapse imaging in (C). **E** Additional representative kymographs, further examples in Supplementary Figure 12DE. Scale as in (D). **F** 2-color time-lapse 2D-SIM imaging *TagF_oex_* sheath (TssB-mCherry2) and TagZ-mNG localization when expressed from the native site and under overexpression conditions, left and right panels respectively. **G** CPRG accumulation assay comparing T6SS-dependent *E. coli* lysis of TagZ-mNG strains to assess complementation. **H** Cartoon visualizing TagZ-mChY topology and tested truncation constructs. **I** CPRG accumulation assay comparing T6SS-dependent *E. col*i lysis of TagF_oex_ phenotype in the TagZ truncation constructs to assess complementation.

TagZ is conserved within TSA loci, yet its function remains unclear, as Δ*tagZ* strain retains T6SS activity (Ringel *et al*, 2017). To address this, we examined TagZ homologs in more detail. MSA analysis of all identified TagZ homologs showed that the proteins share an overall topology with a short cytosolic N-terminus, conserved N-terminal sequence motif S*LRIL***KKK, a single transmembrane-helix, and a variable C-terminal periplasmic domain lacking predicted fold or function (Figure 4B, Supplementary Figure 13A).

Considering the impact of a *tagZ* mutation on TagF_oex_, we asked whether TagZ expression is required to maintain the TagF controlled spike-insertion checkpoint. Interestingly, when we deleted *tagZ* in *ΔvgrG1-4* and compared sheath dynamics in 2D-SIM, we observed ineffective short assemblies for *ΔvgrG1-4ΔtagZ*, albeit at a frequency (0.109 ± 0.017, n = 231 cells) lower than we observed in *ΔvgrG1-4ΔtagF* (Figure 4CDE, Supplementary Figure 12DE).

To understand subcellular localization of *A. baylyi* TagZ, we generated a TagZ C-terminal mNG fusion, expressed it from the native locus (TagZ-mNG), or constitutively overexpressed in *trans* from a neutral site in the genome under rpsL_P_ (TagZ-mNG_oex_). 2D-SIM imaging of TagZ-mNG revealed localization to the cell periphery. TagZ-mNG formed discrete foci, whereas TagZ-mNG_oex_ showed uniform localization (Figure 4F). Surprisingly, T6SS sheath dynamics in the TagZ-mNG overexpressing strain were abrogated. Correspondingly, we observed a significant drop in T6SS dependent lysis of *E. coli* for TagZ_oex_, with no difference in accumulated CPRG absorbance to the T6SS-negative strain *Δhcp* (Figure 4G, Supplementary Figure 13B; *p* = 0.60). Importantly, there is no difference in prey lysis between TagZ-mNG_oex_ and TagZ_oex_ indicating that TagZ-mNG fusion is at least partially functional.

This observation led us to further investigate the functional link between TagZ and TagF. To map the TagZ domains required for TagF function, we generated truncation mutants (Figure 4H). Strains were engineered to express C-terminal mCherry2-tagged TagZ truncations from the native locus. From the native *tagZ* locus, we expressed full-length TagZ-mCherry2 or truncations lacking the C-terminal periplasmic domain (TagZΔperi-mChY) or both the periplasmic domain and transmembrane region (TagZΔperi+TM-mChY). TagF was then overexpressed in each background, and complementation of TagF-dependent T6SS inhibition was assessed by monitoring *E. coli* lysis. We found that full-length TagZ-mChY and the TagZΔperi-mChY truncation both complemented the TagF overexpression phenotype, while TagZΔperi+TM-mChY largely failed to maintain inhibition, producing a partially T6SS-active phenotype similar to TagF_oex_ in the Δ*tagZ* background (Figure 4I, Supplementary Figure 13C).

Taken together, our results show that TagZ localizes to the cell periphery and is essential for TagF function, but the periplasmic domain is dispensable for TagF-mediated T6SS inhibition.

### TagZ N-terminus recruits TagF to the membrane

Exploring a potential direct interaction between TagZ and TagF, we used AlphaFold3 (Abramson *et al*, 2024) to model the complex. The prediction positioned the TagZ N-terminus with high confidence inside the conserved groove at the domain interface of TagF (PAE < 2.5 Å for interacting residues as in Figure 5B, Supplementary Figure 14). This model fit well with our data that indicated that only the first 26 N-terminal residues of TagZ are accessible to cytosolic TagF (Figure 4HI, Supplementary Figure 13A). Furthermore, the model positioned the TagZ residues with the highest conservation among homologs within the TagF groove, where we already identified residues essential for TagF activity (D). Complex interface analyses using PDBePISA (Krissinel & Henrick, 2007) showed an interaction network of one salt bridge and seven hydrogen bonds and total buried surface of 1436 Å^2^ indicating additional contribution of hydrophobic interactions (Figure 5B, Supplementary Figure 14B).

**Figure 5:**
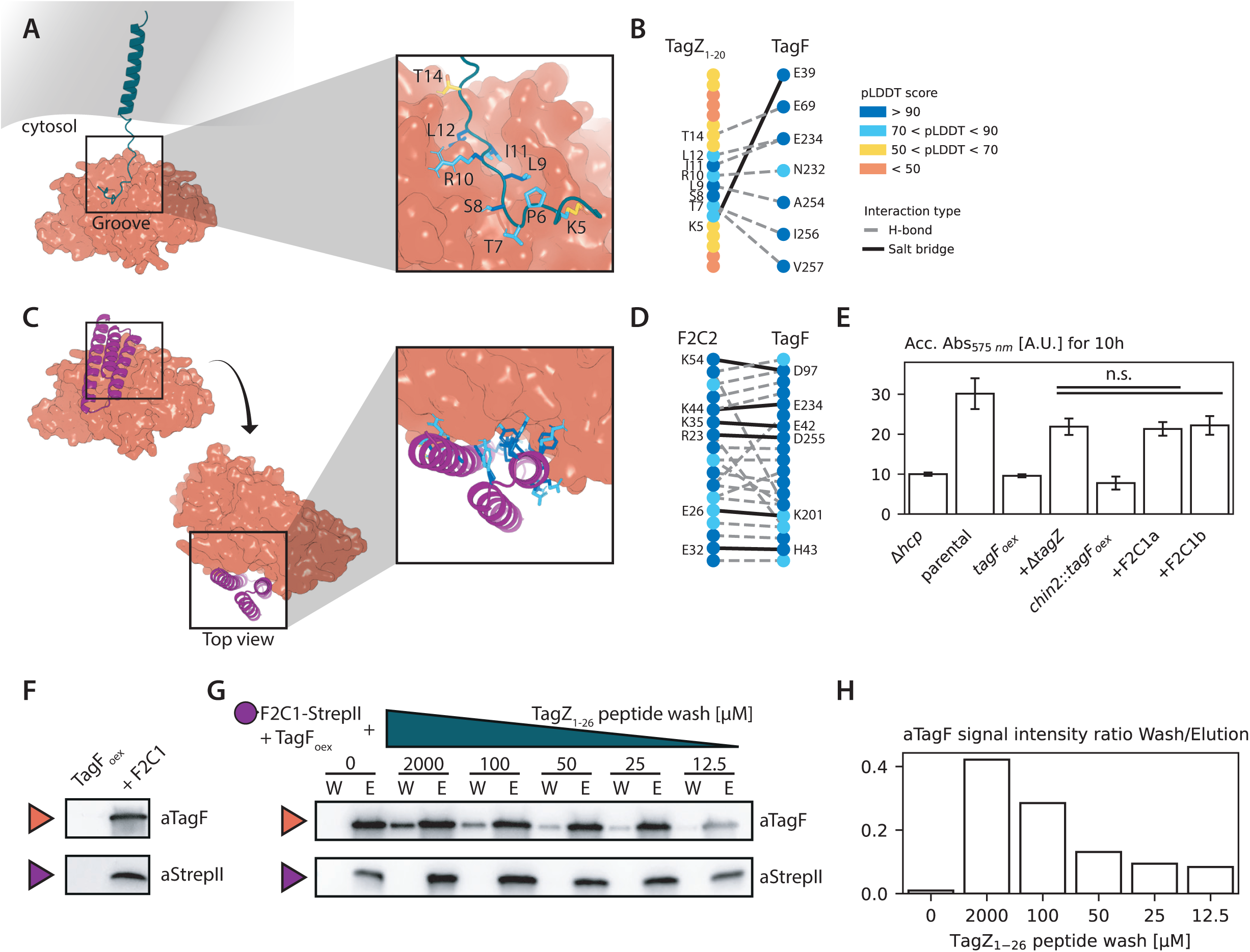
TagF-TagZ direct interaction as suggested by AlphaFold model and demonstrated by showing competition between an occluding binder design and TagZ_1-26_ N-Terminal peptide. **A** TagF (orange) TagZ (teal) AlphaFold3 complex model places the TagZ N-terminus in the TSA-conserved TagF groove. TagZ residues within 4 Å of TagF are shown in stick-representation. AlphaFold per-residue confidence (pLDDT-score) is shown for TagZ interacting residues, full model quality analysis in Supplementary Figure 14A. **B** AlphaFold3 TagF-TagZ interaction network based on PDBePISA analysis. AlphaFold pLDDT-score is shown for interacting residues. **C** TagF-F2C1 binder (purple) AlphaFold3 complex model occluding the TagZ-binding site. F2C1 residues within 4 Å of TagF are shown in stick-representation. AlphaFold pLDDT-score is shown for F2C1 interacting residues, full model quality analysis is shown in Supplementary Figure 15D. **D** AlphaFold3 TagF-F2C1 interaction network based on PBDePISA analysis. AlphaFold pLDDT-score is shown for interacting residues. **E** CPRG accumulation assay comparing T6SS-dependent *E. coli* lysis of TagF_oex_ΔtagZ and TagF_oex_F2C1_oex_. **F** StrepII-tag based co-purification of TagF with F2C1-strepII analyzed by western blotting. **G** StrepII-tag based co-purification of TagF with F2C1-strepII challenged using increasing concentrations of TagZ_1-26_ peptide and analyzed by western blotting. W: TagZ_1-26_ wash, concentrations as indicated; E: 50 mM biotin elution. **H** Relative anti-TagF signal intensity from peptide-wash/biotin-elution ratio in (G) shows a concentration dependent competition effect.

FIDA binding experiments with purified TagF and the soluble TagZ_1-26_ N-terminal peptide only showed weak binding in isolation, suggesting a low-affinity transient interaction or a requirement for cellular context to increase affinity (Supplementary Figure 14D). We therefore focused on a cellular approach by de-novo designing a protein that would bind the TagF groove to block the putative binding site of TagZ.

For the design of site-specific TagF-binding proteins, we used RFdiffusion (Watson *et al*, 2023) for protein structure design in combination with ProteinMPNN (Dauparas *et al*, 2022) for protein sequence design. We then used AlphaFold2 (Jumper *et al*, 2021) and Rosetta (Leaver-Fay *et al*, 2011) to evaluate the designed amino acid sequences and selected 24 candidates for experimental screening in *A. baylyi* (Supplementary Figure 15A). Co-expression of TagF and each binder in parallel from neutral genomic sites revealed that binder F2C1 effectively blocked TagF function using a lysis assay (Figure 5CE, Supplementary Figure 15BCD). Compared to TagZ, PDBePISA analysis of the TagF-F2C1 complex showed an extended hydrogen bonding network with several key salt bridges stabilizing the interaction (Figure 5D, Supplementary Figure 15E). The increased surface area of 2729 Å^2^ could therefore effectively block TagZ binding. Co-purification of TagF with F2C1-strepII from *A. baylyi* lysate confirmed that the effect on TagF function resulted from a direct interaction (Figure 5F, Supplementary Figure 16A).

We then tested if the binder occupies the groove on TagF as predicted for TagZ. We ran parallel co-purifications as described above, this time washing immobilized F2C1-strepII and enriched with TagF using increasing concentrations of TagZ_1–26_ peptide prior F2C1 to elution from the beads. Importantly, addition of TagZ_1–26_ peptide led to a concentration-dependent elution of TagF from F2C1, consistent with direct binding at the same site and competitive displacement (Figure 5GH, Supplementary Figure 16BC).

Based on these results, we conclude that the N-terminus of membrane localized TagZ interacts with TagF groove. This would result in a preferential localization of TagF to the cell periphery and thus increase its local concentration at T6SS assembly initiation sites where it would allow assembly of the full length T6SS sheath upon spike incorporation to the baseplate.

## Discussion

Here we show that in *Acinetobacter*, T6SS assembly progresses through a TagF-controlled regulatory checkpoint which prevents sheath-tube assembly in the absence of an assembled spike complex and therefore ensures successful secretion of T6SS substrates (Figure 6).

**Figure 6:**
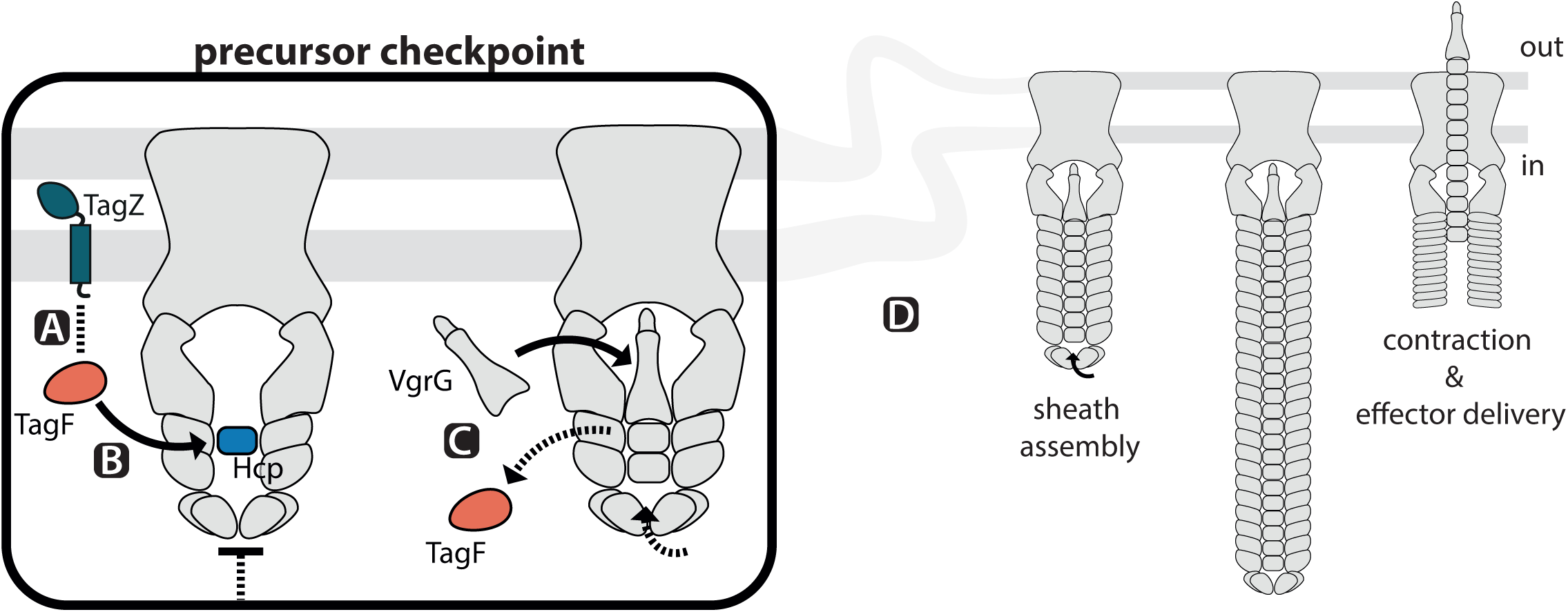
Proposed model of the precursor checkpoint mechanism. **A** Direct but transient TagF-TagZ interaction increases local concentration of TagF near the inner membrane. **B** TagF prevents T6SS sheath assembly by interfering with Hcp ring formation at the pre-cursor. **C** VrgG-spike displaces TagF at the precursor. **D** Once VgrG/PAAR is positioned within the baseplate, Hcp tube formation and sheath assembly resume and eventually result in successful sheath contraction and effector delivery.

TagF is a conserved post-translational inhibitor of T6SS dynamics that was described in regulatory context with Fha and the TPP kinase-phosphatase pair PpkA/PppA , where upon a specific signal, Fha is phosphorylated by PpkA and promotes T6SS assembly (Mougous *et al*, 2007; Boyer *et al*, 2009; Hsu *et al*, 2009; Silverman *et al*, 2011). However, TagF homologs were also identified in species lacking Fha as the putative TagF interaction partner, and without a dedicated phosphatase-kinase pair (Bayer-Santos *et al*, 2019; Boyer *et al*, 2009; Schwarz *et al*, 2010; Weber *et al*, 2016). Our bioinformatic analyses showed that 22% of *tagF* genes were encoded in T6SS clusters lacking *fha* and the phospho-regulatory TPP relay. We identified a group of TagF proteins, that were co-encoded with an alternative set of conserved accessory genes, T6SS Accessory Gene Set A (TSA), which included *tsmK*, *tagN*, *tagX*, *tslA*, and *tagZ*.

In *P. aeruginosa*, PaTagF1 was shown to be a key post-translational regulator of T6SS dynamics, where it blocks most assemblies at native expression levels (Basler *et al*, 2013; Silverman *et al*, 2011). Overexpression of TagF homologs in *A. tumefaciens* and *S. marcescens* was previously shown to block T6SS dynamics, suggesting an analogous regulatory mechanism (Lin *et al*, 2018; Ostrowski *et al*, 2018). We show that overexpression of *A. baylyi* TagF leads to inhibition of T6SS dynamics and formation of stable sheath foci localized to the cell periphery, suggesting that TagF blocks assembly at a defined assembly checkpoint (Figure 1F). Previously, similar structures were identified in an *A. baylyi ΔtagX* mutant, where such T6SS precursors were preferentially localized to cell-cell contact sites by the accessory protein TslA (Lin *et al*, 2022). We also observed the accumulation of TssA foci at cell-cell contacts upon TagF overexpression, suggesting that TslA-dependent localization of precursors precedes the TagF-mediated block of sheath elongation. Indeed, TagF is not required for precursor positioning, as we also observed precursor formation at cell-cell contacts in TagF deletion strains (Supplementary Figure 5C).

Integration of the VgrG/PAAR spike loaded with effectors into the assembly is required for T6SS function. Previously, T6SS assembly has been repeatedly linked to VgrG/PAAR and effector loading in different species, including *Acinetobacter* (Shneider *et al*, 2013; Liang *et al*, 2015; Lopez *et al*, 2020; Wu *et al*, 2020; Donato *et al*, 2020). In agreement with these reports, we observed that deletion of all four *A. baylyi vgrG* genes resulted in a loss of T6SS dynamics, where only sheath precursors remained. Surprisingly, deletion of *tagF* in the *ΔvgrG1-4* background partially restored sheath dynamics suggesting that removing the TagF-mediated regulatory checkpoint allows assemblies lacking the spike and effectors. Collectively, this suggests that TagF is actively blocking sheath-tube assembly until the spike is loaded to the baseplate, and that spike-integration is not necessary to trigger sheath elongation.

Next, we sought to uncover structural differences between homologs in T6SS clusters encoding TagF in TPP or TSA contexts. In *P. aeruginosa* PaTagF1 the Fha1 binding interface is specifically formed by two N-terminal surface-exposed loops α and β (Lin *et al*, 2018). Comparing the available structure of PaTagF1 to the *A. baylyi* TagF crystal structure, we found two loops in the N-terminal domain with high structural similarity to the Fha1-interaction site on PaTagF1, which were also essential for TagF-mediated arrest of T6SS dynamics in *A. baylyi* (Filippova *et al*, 2007; Lin *et al*, 2018). This suggests that the N-terminal domain is a structurally conserved binding site. However, *P. aeruginosa* and *A. baylyi* T6SS assemblies are distinct, particularly at the membrane-complex baseplate interface. *P. aeruginosa* H1-T6SS encodes both for TssJ1 and Fha1, whereas *Acinetobacter* T6SS instead requires TsmK as an additional part of membrane complex (Chen *et al*, 2015; Kandolo *et al*, 2023; Ringel *et al*, 2017). These differences in overall T6SS assembly are also reflected in the distinct architectures of TagF proteins, as we identify a C-terminal domain in *A. baylyi* TagF that is not present in PaTagF1, which contributes to a new interaction surface on *A. baylyi* TagF formed by a groove at the interface between its two domains.

The differences in T6SS assembly are also carried over to direct interaction partners. While PaTagF1 binds Fha1, we show that *A. baylyi* TagF instead interacts with Hcp. Importantly, we show the observed hydrodynamic radius of TagF-Hcp complex was smaller than a full hexameric Hcp_6_ ring, giving strong indication that TagF blocks T6SS assembly by actively breaking the ring structure (Figure 3F). This supports the model where during assembly, TagF is displaced from Hcp by the spike to allow Hcp_6_ ring formation and thus sheath-tube polymerization. Such a mechanism would ensure that the VgrG/PAAR/effector complex is properly inserted into the baseplate before the energetically expensive step of sheath polymerization, contraction and disassembly is initiated and prevents that the system fires ‘blanks’, i.e. empty tube. This mirrors findings that the presence of effectors is required for T6SS activity in *V. cholerae and A. tumefaciens* (Liang *et al*, 2019; Wu *et al*, 2020). Moreover, our findings parallel the hierarchical secretion control observed in Type III Secretion Systems (T3SS). In *E. coli* (EPEC), the regulatory SctW gatekeeper complex associates with the translocase gate SctV, promoting middle-substrate/chaperone interactions while blocking effector/chaperone loading but is dislodged upon host-contact to facilitate effector translocation (Portaliou *et al*, 2017; Díaz-Guerrero *et al*, 2021; Armentrout & Rietsch, 2016).

Measured dissociation constant for TagF-Hcp complex indicates that at the native expression levels, TagF cannot fully saturate all free Hcp in the cytosol (Lin *et al*, 2019a). Consequently, the observed block on T6SS dynamics would require recruitment of TagF to the T6SS assembly site to monitor spike insertion. We demonstrate that the accessory protein TagZ, which is conserved in 85% of the TSA regulatory modules we analyzed, is required for effective TagF-mediated regulation of T6SS. Deletion of *tagZ* in a strain overexpressing TagF resulted in a loss of TagF-mediated inhibition of T6SS activity. Moreover, deletion of *tagZ* in *ΔvgrG1-4* strain resulted in dynamic, short assemblies similar to those observed upon *tagF* deletion in the same background (Figure 4AC). TagZ is predicted to be a single-pass membrane protein with a disordered periplasmic domain and only a short N-terminal peptide exposed to the cytosol. AlphaFold-based modeling suggested that this peptide could interact with the conserved groove on *A. baylyi* TagF, and using FIDA, we did indeed observe weak binding (Figure 5A, Supplementary Figure 14D). Importantly, overexpression of F2C1, a synthetic binder targeting the conserved TagF groove, disrupted TagF function, while *in vitro*, the TagZ N-terminal peptide competed for the same site and could displace F2C1 (Figure 5). These data support a model in which membrane anchored TagZ recruits TagF through its cytosolic peptide. This would increase local concentration of TagF close to the site of T6SS baseplate assembly and thus prevent Hcp_6_ ring formation and sheath-tube elongation until the spike is loaded. Moreover, TagZ may coordinate additional assembly steps in response to membrane complex formation, localization, or alternative periplasmic cues.

Together, our results greatly expand the scope of known T6SS regulatory genes and show that TagF is a checkpoint regulator whose function goes beyond simple control of T6SS assembly frequency. We show that TagF is responsible for coordinating spike positioning within the baseplate prior to full assembly. Spike-assembly and effector-loading has now been repeatedly identified as central for function in several T6SS and may present a shared checkpoint for different regulatory networks (Liang *et al*, 2015; Lopez *et al*, 2020; Shneider *et al*, 2013; Wu *et al*, 2020). TagF interactions with conserved T6SS proteins imply that part of the regulatory mechanism may be conserved over a divergent collection of T6SS variants. Collectively, our results stress the complexity of T6SS regulation coordinating assembly to ensure effective secretion. We anticipate that other T6SS may utilize parallel regulatory pathways that also target the early steps of assembly.

## Materials and Methods

### Bioinformatic analyses

#### Genomic context analyses

GC analysis was performed using GCSnap, the tool kit is available under GPL3.0 license (Pereira, 2021). As input for GCSnap, TagF homolog protein coding sequences were filtered to 70% maximum identity. GCSnap analysis was performed including 16 genes flanking the target sequence on either side. GCSnap output is available in json file format in the extended data. To restrict our downstream analysis to functional T6SS loci, we filtered the GCCnap output to loci encoding at least three out of four homologs of a subset of essential T6SS genes (TssM, Hcp, TssC, and ClpV). Key input and output files used during analysis are provided in Supplementary File 1.

#### Sequence alignments

MSA protein sequence alignment and conservation analyses were performed in jalview using the Clustal Omega algorithm (Sievers *et al*, 2011; Waterhouse *et al*, 2009). Key input and output files used during analysis are provided in Supplementary File 3.

#### Evolutionary analysis/ Phylogenetic tree

Amino acid sequence alignments were performed on a set of 655 amino acid sequences retrieved from NCBI based on the TagF-sequences used in GCSnap analyses. The alignment was performed using the MUSCLE algorithm (Edgar, 2004). Phylogeny was inferred using the Maximum Likelihood method and Jones-Taylor-Thornton model (Jones *et al*, 1992) of amino acid substitutions and the tree with the highest log likelihood (-80,413.40) was used for further analysis. The percentage of replicate trees in which the associated taxa clustered together, where the number of replicates (112) was determined adaptively (Kumar *et al*, 2024). The initial tree for the heuristic search was selected by choosing the tree with the superior log-likelihood between a Neighbor-Joining (NJ) (N Saitou & M Nei, 1987) tree and a Maximum Parsimony (MP) tree. The NJ tree was generated using a matrix of pairwise distances computed using the Jones-Taylor-Thornton model (Jones *et al*, 1992). The MP tree had the shortest length among 10 MP tree searches; each performed with a randomly generated starting tree. The partial deletion option was applied to eliminate all positions with less than 95% site coverage resulting in a final data set compromising 131 positions. Evolutionary analyses were conducted in MEGA12 utilizing up to 18 parallel computing threads (Kumar *et al*, 2024). Tree visualizations and annotations were performed using iTOL (Letunic & Bork, 2024). The input data used for tree computation, MEGA12 output, and the full annotated tree are available in Supplementary File 2.

### Strains and growth conditions

*A. baylyi*, *E. coli*, and *P. aeruginosa* strains were cultivated in lysogeny broth (LB) liquid media or LB-agar plates supplemented with 1.5% agar. *A. baylyi* was cultivated at 30°C, *E. coli* was cultivated at 37°C, unless stated otherwise. For *A. baylyi* cultivation, antibiotic concentrations were 50 µg/ml kanamycin and 50 µg/ml streptomycin as required for selections. *E. coli* cultures were supplemented with 50 µg/ml kanamycin, 100 µg/ml ampicillin, 50 ug/ml chloramphenicol, or 15 µg/ml gentamicin, dependent on the respective selection resistance. Day cultures were prepared free of antibiotics, apart from *E. coli* overexpression cultures. Strains were archived at -80°C, stocks were prepared from overnight culture and mixed with 16% dimethyl sulfoxide (DMSO) as cryoprotectant. A full list of all strains used in this study is summarized in Supplementary Table 1.

### Construction of *A. baylyi* chromosomal mutants

*A. baylyi* chromosomal mutants were created as previously described (Metzgar *et al*, 2004), with homologous regions for integration measuring between 450 and 700 bp. Chromosomal overexpression strains were constructed by cloning the endogenous strong *rpsL* ribosomal promoter in front of the gene of interest, and an additional downstream kanamycin resistance marker for selections. Overexpression cassettes were created with homologous regions for integration into the *ampC* gene, a neutral location previously used in (Lin *et al*, 2022), and an additional integration site chin2 located in the intergenic region between *ACIAD2126* and *ACIAD2129*. Oligonucleotides used to generate the various constructs are summarized in Supplementary Table 2.

### Construction of *E. coli* heterologous expression strains

For large scale protein overexpression intended for purifications, genes of interest were cloned into expression vectors using the FX-cloning platform (Geertsma, 2013). In a first step, genes were amplified by PCR with asymmetric SapI restriction site overhangs (designed on fxcloning.org) and cloned into pINIT.cat via restriction-ligation. Chemically competent *E. coli* MC1061 cells were transformed with FX-ligation mixes and plated on LB-agar supplemented with antibiotic for selection. Plasmids were isolated using PureYield Plasmid miniprep kit (Promega) and sequenced using the Sanger method (Microsynth). After sequence confirmation, genes in pINIT.cat were subcloned into the pBXC3H expression vector (Addgene). After transformation of *E. coli* MC1061 and selection on plate, sequenced clones were archived and used for overexpression cultures. A full list of all plasmids created and used in this study is summarized in Supplementary Table 3.

### CPRG lysis assay

*A. E. coli* lysis assays were performed based on (Ringel *et al*, 2017). Biological triplicate cultures of *A. baylyi* strains of interest and the *E. coli* MG1655 *lacZ+* prey strain B252 were inoculated in LB supplemented with appropriate antibiotics and cultivated overnight. The following day, day cultures of all strains were inoculated from liquid overnight culture and cultivated for approximately 3 h until *A. baylyi* cultures reached OD_600_ = 0.8-1.2. *E. coli* cultures were supplemented with 0.1 mM isopropyl-beta-D-1-thiogalactopyranoside (IPTG) to induce LacZ expression already during cultivation. All cultures were concentrated to OD_600_ = 10. Killing mixes were prepared in technical triplicate with a 1:5 ratio of predator to prey, typically in volumes of 20 µl : 100 µl. Using a pin tool (VD380D from V&P SCIENTIFIC Inc.), 1.5 µl of all mixes were transferred to single-well plates (Nunc® OmniTray, MaxisorpTM) filled with 30 ml of LB-agarose supplemented with 0.1 mM IPTG and 20 µg/ml Chlorphenolred-β-D-galactopyranosid (CPRG). The LacZ-dependent conversion of CPRG to chlorophenol-red was followed by monitoring absorbance at 575 nm every 5 minutes at 30°C, for up to 20 h using an EPOCH2 microplate reader (Agilent BioTek). For data analysis, initial reads at time point zero were averaged over the entire plate and used to blank all following measurements. Lysis accumulation was calculated by forming the sum of all absorbance values recorded per biological replicate in the first 10 h. Statistical significance was determined using the ANOVA method (* *p* < 0.05, ** *p* < 0.01, *** *p* < 0.001, **** *p* < 0.0001).

### Structured Illumination Microscopy (SIM) of *A. baylyi*

*A. baylyi* cells were restreaked from overnight LB-agar culture supplemented with appropriate antibiotics on fresh LB-agar and cultivated for 2 h at 30°C. Thin LP agarose pads (1:2 (v/v) LB to phosphate buffered saline (PBS), 1% agarose, 0.25-0.4 mm thick) were freshly prepared and cut into equal squares of approximately 4 mm^2^. Cells were scraped off the agar and resuspended in LB at OD_600_ = 15, mixing thoroughly to dissolve clumps. 2.5 µl of cell suspension was spotted onto each agar pad and a clean coverslip (1.5H from Paul Marienfeld GmbH) was placed on top. Data acquisition for optimal dynamics and reproducibility was performed between 5 and 20 minutes after specimen preparation. SIM imaging was performed on a Deltavision OMX SR equipped with an Olympus 60x PLAPON objective lens (1.412A), UltimateFocus Hardware Autofocus module with Focus Assist, and sCMOS cameras. 2D-SIM time lapses of sheath dynamics (TssB-msfGFP) were recorded at a frame rate of 5 s for up to 3 minutes using a 488 nm laser with an exposure of 10 ms and a laser power of 10%, resulting in approx. 2500-5000 counts on the initial frames. TssA-mNG and TagZ-mNG imaging was recorded using a higher exposure of 25 ms and an increased laser power of 20%. TssB-mCherry2 dynamics were recorded using a 603 nm laser using 20% laser power and 25 ms exposure. SIM reconstructions were performed using DeltaVision OMX softWoRx. Pixel size of the reconstructed images was 0.08 x 0.08 x 0.38 µm (x y z).

### Image analysis

Image analysis was performed in Fiji (Schindelin *et al*, 2012). Due to the intensity distribution from SIM reconstruction, auto-segmentation performance was poor and quantifications were instead performed manually. Imaging data from three separate biological replicates (either different clones or the same clone imaged on different days) and at least two different field of views each were analyzed for each strain. For manual quantification, 2 to 3 sub-areas of 300px^2^ were selected per FOV. **Cell counts** were performed on the initial frame of the timelapse, all cells that were considered for T6SS-related quantifications were included in the cell count (cells fully in frame and cells where at least half the volume was in frame). **Sheath assembly frequencies** were determined following full T6SS sheath assembly and contraction events throughout the timelapse, as assemblies can occur multiple times at the same location. Only assemblies that originated from within the focal plane, directly at the cell periphery, were quantified. Where applicable, statistical significance was determined using the ANOVA method (* *p* < 0.05, ** *p* < 0.01, *** *p* < 0.001, **** *p* < 0.0001).

For the **maximum sheath structure length analysis**, timelapses were aligned using XTurboStackReg (Thevenaz *et al*, 1998) and then a projection of the sum intensity of all frames was performed prior to structure length measurements. Only assemblies that originated from within the focal plane, directly at the cell periphery, were considered for the measurements. Note that we chose to use the sum of all frames approach rather than maximum intensity projections: Assemblies appear shorter, as the most extended state typically only occurs on 1/37 frames of a 3 min timelapse. While maximum intensity projections gave better results for lower frequency T6SS assemblies such as Δ*vgrG1-4ΔtagF* (capturing full assembly length), they were less reliable for the parental strain due to increased sheath dynamics. Both approaches yielded similar overall trends in the data. For manual quantification, per FOV, 3 areas of 300px^2^ were selected for manual quantification. As the data distribution was determined to be non-normal in the case of the maximum structure length analysis, significance was determined using Dunns’ multiple comparisons test and Kruskal-Wallis-ANOVA test.

### Heterologous protein expression for purifications

Proteins TagF-His_10_ and FLAG-Hcp were produced in *E. c*oli MC1061 with a 3C-protease cleavable affinity tag from the arabinose-inducible plasmid pBXC3H (Addgene,(Geertsma, 2013)). For small scale co-purifications, His-MPB, and FLAG-MBP were produced similarly but with non-cleavable N-terminal affinity-tag fusions. For protein purifications, overexpression cultures were prepared with LB supplemented with antibiotic, with a 1% inoculum from liquid overnight pre-culture. Expression cultures were cultivated at 37°C, 200 rpm until OD_600_ = 0.4, after which the temperature was reduced to 25°C for 30 min. Expression was then induced through addition of 0.005% (w/v) L-arabinose. The overexpression cultures were harvested 12 to 14 h post-induction by centrifugation at 5,500 g for 10 min at 4°C. Harvested material was stored at -20°C until purification.

### Small-scale co-purifications for interaction analyses

Frozen cell pellets were thawed on ice and resuspended in ice-cold purification buffer (20 mM Hepes pH 7.5, 150 mM NaCl, 10% glycerol) to OD600 = 200. Lysozyme (1 mg/ml), DNaseI (1 ug/ml), MgSO4 (1 mM) and cOmplete EDTA-free protease inhibitor (ROCHE) were added as lysis supplements. After 1 h of incubation at 4°C, mixing, cells were lysed by sonication using an Eppendorf-tube sonicator in 4 cycles of 30 s on 60 s off (75% power, 50% on-time). Lysate was clarified by centrifugation for 10 min at 16,000 g at 4°C. Specific to the respective affinity tag targeted, 50 µl bead slurry per sample was equilibrated in batch by washing with 1 ml each of purification buffer, blocking buffer (purification buffer supplemented with 0.5% bovine serum albumin (BSA) and three more times in purification buffer prior to sample application. The specific beads used in the different co-purifications, as well as the appropriate washing and elution schemes are detailed below. Eluted samples were stored at -20°C until processing for mass spectrometry or western blotting analysis.

#### Anti-His_10_-tag co-purifications

Equilibrated HisPur™ Ni-NTA Magnetic Beads (Thermo Fisher) were loaded with 150 to 300 µg of purified bait protein or 1 ml of clarified cell lysate in presence of 15 mM imidazole (minimum 4x bead binding capacity). Samples were incubated 30 min at 4°C, mixing end over end. Post-binding, beads were washed three times with 750 µl washing buffer (purification buffer supplemented with 75 mM imidazole). Subsequently, 1 ml clarified prey lysate was added to each sample and incubated for 30 min at 4°C, mixing. For the initial screen for TagF-interacting proteins, *A. baylyi* lysate solubilized in 1.6% DDM (n-dodecyl-beta-D-maltoside, non-ionic detergent) and clarified by ultracentrifugation was used to include membrane proteins in the screen. For more focused experiments later-on, DDM-solubilization was omitted. Afterwards, beads were washed three times with 750 µl washing buffer (supplemented with 75 mM imidazole). For the elution, beads were resuspended in 100 µl purification buffer supplemented with 300 mM imidazole.

#### Anti-FLAG-tag co-purifications

Equilibrated ANTI-FLAG® M2 magnetic beads (Merck) were loaded with 1 ml of clarified cell lysate. Samples were incubated 30 min at 4°C, mixing end over end. Post-binding, beads were washed three times with 750 µl purification buffer. Subsequently, 1 ml lysate containing prey protein was added and incubated for 30 min at 4°C, mixing end over end. Beads were washed three times with 750 µl purification buffer. For the elution, beads were resuspended in 100 µl purification buffer supplemented with 0.6 mg/ml FLAG peptide (GenScript).

#### Anti-strepII-tag co-purification of binder

Equilibrated MagStrep Strep-TactinXT beads (Fisher Scientific) were loaded with 1 ml of clarified cell lysate. Samples were incubated 30 min at 4°C, mixing end over end. Post-binding, beads were washed three times with 750 µl purification buffer. For the **TagZ-peptide competition experiment**, a two-fold dilution series of TagZ_1-26_ peptide (2 mM to 12.5 µM) in purification buffer was incubated with the loaded beads for 30 min at 4°C, mixing end over end. Here, two additional washes with 750µl purification buffer were performed afterwards. For the elution, beads were resuspended in 100 µl purification buffer supplemented with 50 mM biotin.

#### Large-Scale Purifications for biophysical and structural characterizations

Frozen cell pellets were thawed at room temperature and resuspended in ice-cold purification buffer (20 mM Hepes pH 7.5, 150 mM NaCl, 10% glycerol) to OD600 = 200. Lysozyme (1 mg/ml), DNaseI (1 ug/ml), MgSO4 (1 mM) and cOmplete EDTA-free protease inhibitor (ROCHE) were added as lysis supplements. After 1 h of incubation at 4°C, mixing, cells were lysed by sonication for a total of 5 min at 50% power (Branson Sonifier SFX550). Lysate was clarified by centrifugation for 30 min at 35,000 rpm (Beckman Ti45 rotor), 4°C. The supernatant was collected and passed through a 0.450 µm syringe filter. Lysate was then purified using the specific affinity-tag fusions using applicable affinity-chromatography resin (specified below). Pooled elution fractions were dialyzed at 4°C for a minimum of 1 h or overnight using 6-8 molecular weight cut-off (MWCO) dialysis tubing (SpectraPor®) against purification buffer. The affinity tag was routinely cleaved by addition of 0.2 mg/ml 3C-protease during dialysis, unless it was specifically required for experiments. Protein recovered from dialysis was concentrated up to 25 mg/ml using a molecular weight cutoff filter of appropriate dimensions (Merck Amicon). Samples were further purified by size exclusion chromatography (SEC) in 20 mM Hepes pH 7.5, 150 mM NaCl using an Äkta Pure 25 chromatography system (Cytiva) using a HiLoad 75 16/600 Superdex pg column for TagF and a Superdex 200 increase 10/300 GL for Hcp. SEC peak fractions were pooled and concentrated as required for experiments. Samples taken during all steps of purification were routinely analyzed on SDS-PAGE and via western blotting. Protein not used immediately for experiments was aliqoutted, flash-frozen in liquid nitrogen, and stored at -80°C.

#### TagF IMAC-purification step

Immobilized metal affinity chromatography (IMAC) was performed using an Äkta Start (Cytiva) peristaltic pump and a column packed with His-Fastflow Sepharose Ni-nitrilotriacetic acid (NTA), pre-equilibrated with purification buffer. The filtered lysate was loaded using the sample pump at a flow rate of 1 ml/min and the column was washed with 5 column volumes (CVs) purification buffer. An additional washing step was performed with 10 CV purification buffer supplemented with 50 mM imidazole. Protein of interest was eluted in 5 CV at 0.5 CV fractionation, using 300 mM imidazole in purification buffer. Pooled elution fractions were immediately dialyzed to decrease imidazole concentrations below 10 mM, we found TagF to be sensitive to prolonged exposure to higher concentrations of imidazole.

#### Hcp FLAG-purification step

FLAG affinity purification was performed using 6 ml Anti-DYKDDDDK G1 Affinity Resin (GenScript) in a gravity column. Resin was washed with 20 CV PBS and equilibrated using 20 CV purification buffer. Batch binding with lysate was performed for 1 h at 4°C, agitating. Afterwards, the resin was returned to the column, drained and washed twice with 20 CV purification buffer. FLAG-Hcp was eluted from the resin using 1 CV of purification buffer with 0.6 mg/ml FLAG peptide (GenScript), and an additional three washing steps of 1 CV of purification buffer with 0.1 mg/ml FLAG peptide were used to improve yields.

### SDS-PAGE and western blotting

Protein samples were mixed with 4X Laemmli protein sample buffer (BioRad) and applied to 4-20% Mini-PROTEAN TGX Precast protein gels (Biorad) for Tris-Glycine SDS-PAGE. Gels were routinely stained with QuickBlue Protein Stain (LuBioScience). For immunoblotting, proteins were transferred from gel to a nitrocellulose membrane (Cytiva) using a Trans-Blot Turbo Transfer System (Biorad) for semi-dry western blotting with transfer buffer (48 mM Tris, 39 mM glycine, 20% methanol (v/v)). Blotted membranes were blocked for 1 h or overnight in 5% (w/v) skim milk powder in PBST (PBS+ 1% Tween20). All antibodies were diluted in 5% (w/v) skim milk powder in PBST. Blots were washed 3 times for 5 min in PBST between different antibody treatments and prior to blot development. All antibodies used in this study are summarized in Supplementary Table 4. Secondary antibodies were coupled to horse radish peroxidase. Blots were developed using the LumiGLO Peroxidase Chemiluminescence Kit (SeraCare) and documented using a Fusion FX7 Edge imaging system (Witec AG).

### TagF crystallization

Purified TagF in 20 mM Hepes pH7.5, 150 mM NaCl buffer was concentrated to 15 mg/ml using a 30 kDa MWCO filter (Merck Amicon). Screening of TagF crystallization conditions identified one suitable condition with sitting drop vapor diffusion in a 1:1 sample to buffer ratio yielded crystals in 0.05 M CdSO4 (Salt), 0.1 M Hepes 7.5 pH (Buffer), 1 M NaOAc (Precipitant) after three to four days at 20°C. Additional optimization of the crystallization condition was performed, and crystals were obtained in multiple conditions at both 20°C and 4°C. For X-ray diffraction data collection, crystals were harvested after 3 weeks. Crystals were cryo-protected in mother liquor supplemented with 20% ethylene glycol and frozen in liquid nitrogen. The reservoir of the optimization condition resulting in the highest resolution diffraction data set collected contained 0.162 M CdSO4 (Salt), 0.1 M Hepes 7.5 pH (Buffer), 0.5 M NaOAc pH 8 (Precipitant).

### X-ray diffraction Data collection

Datasets of well diffracting crystals were collected at the Swiss Light Source (Villingen) at the PXIII X06DA beamline on a PILATUS 2M-F detector at X-ray wavelength l = 1.000009 Å. The best crystal diffracted X-rays up to a resolution of 1.8 Å in space group P21P21P21. The asymmetric unit co-ordinated a TagF monomer with cadmium ions from the crystallization condition. The unit cell dimensions were a = 45.1, b = 74.9, c = 85.5, and α = 90°, β = 90°, γ = 90°. Single wavelength anomalous diffraction (SAD) phasing datasets based on the anomalous signal from cadmium ions in the crystal were recorded at 5,700 eV, with the maximum resolution limited to 2.6 Å due to technical constraints (Wang, 1985). Diffraction data was reduced using Dials, scaling and merging was performed with AIMLESS (Agirre *et al*, 2023; Winter *et al*, 2022).

### TagF crystal structure determination

The dataset was experimentally SAD phased with CRANK2 (Skubák *et al*, 2018). The initial model was built using Buccaneer and the remaining residues were manually built in COOT (Cowtan, 2006; Emsley *et al*, 2010). Early refinements were performed in Refmac5 (Agirre *et al*, 2023; Murshudov *et al*, 2011), later in Phenix.refine (Liebschner *et al*, 2019). Structure figures were prepared in PyMOL (Schrödinger & DeLano, 2020). Sequence conservation visualization on the structure was calculated using Consurf server (Ashkenazy *et al*, 2016; Yariv *et al*, 2023) based on provided multiple sequence alignments.

Data and refinement statistics are summarized in Table 1.

**Table 1:** TagF Xray-data and refinement statistics.

| Property | Value | Source |
| --- | --- | --- |
| Space group | P 21 21 21 | Depositor |
| Cell constants<br>a, b, c, $\alpha$, $\beta$, $\gamma$ | 45.15 Å 74.91 Å 85.50 Å<br>90.00° 90.00° 90.00° | Depositor |
| Resolution (Å) Depositor | 56.34 – 1.80<br>56.34 – 1.80 | Depositor<br>EDS |
| % Data completeness<br>(in resolution range) | 100.0 (56.34-1.80)<br>100.0 (56.34-1.80) | Depositor<br>EDS |
| Rmerge | 0.12 | Depositor |
| $\langle I/\sigma(I) \rangle$ | 1.15 (at 1.80 Å) | Xtrriage |
| Refinement program | PHENIX (2.0_5716) | Depositor |
| R, R<sub>free</sub> | 0.209, 0.245<br>0.216, 0.251 | Depositor<br>DCC |
| Wilson B-factor (Å<sup>2</sup>) | 30.6 | Xtrriage |
| Anisotropy | 0.399 | Xtrriage |
| Bulk solvent k<sub>sol</sub>(e/Å<sup>3</sup>), B<sub>sol</sub>(Å<sup>2</sup>) | 0.45,45.1 | EDS |
| L-test for twinning<sup>2</sup> | $\langle \vert L\vert \rangle = 0.48$, $\langle L_2 \rangle = 0.32$ | Xtrriage |
| Estimated twinning fraction | No twinning to report. | Xtrriage |
| F<sub>o</sub>,F<sub>c</sub> correlation | 0.95 | EDS |
| Total number of atoms | 5335 | wwPDB-VP |
| Average B, all atoms (Å<sup>2</sup>) | 47.0 | wwPDB-VP |

<sup>1</sup> Intensities estimated from amplitudes.
<sup>2</sup> Theoretical values of $\langle |L| \rangle$ , $\langle L_2 \rangle$ for acentric reflections are 0.5, 0.333 respectively for untwinned datasets, and 0.375, 0.2 for perfectly twinned datasets.

### Mass spectrometry analyses

#### Pre-treatment for whole cell samples

Bacteria were lysed in buffer containing 5% SDS, 100 mM triethylammonium bicarbonate (TEAB), and 10 mM tris(2-carboxyethyl)phosphine hydrochloride (TCEP) and heated at 95°C for 10 min. Lysates were further sonicated (20 cycles, Bioruptor, Diagnode, 4°C). Carbamidomethylation of cysteins was performed by addition of 20 mM idoacetamide at 25°C in the dark for 30 minutes.

#### Pre-treatment for protein samples

Samples were supplemented with 5% SDS, 100 mM TEAB, 10 mM TCEP and 20 mM iodoacetamide. Samples were incubated protected from light at 25°C for 30 min, shaking at 475 rpm.

#### Sample preparation for LC-MS-based proteome analysis

Proteins were purified and digested with the SP3 approach (Hughes *et al*, 2019) using a Freedom Evo 100 liquid handling platform (Tecan Group Ltd., Männedorf, Switzerland). In brief, Speed BeadsTM (#45152105050250 and #65152105050250, GE Healthcare) were mixed 1:1, rinsed with water and diluted to the 8 µg/µl stock solution. Samples were adjusted to the final volume of 90 µl and 10 µl of the beads stock solution was added to them. Proteins were bound to the beads by addition of 100 µl of 100% acetonitrile to the samples, which were then incubated for 8 min at RT with a gentle agitation (200 rpm). After, samples were placed on a magnetic rack and incubated for 5 minutes. Supernatants were removed and discarded. The beads were washed twice with 160 µl of 70% (v/v) ethanol and once with 160 of 100% acetonitrile. Samples were placed off the magnetic rack and 50 µL of digestion mix (10 ng/µl of trypsin in 50 mM triethylammonium bicarbonate) was added to them. Digestion was allowed to proceed for 12 h at 37°C. After digestion samples were placed back on the magnetic and incubated for 5 minutes. Supernatants containing peptides were collected and dried under vacuum.

#### Mass spectrometry analysis

Sample injection order was block randomized based on the experimental conditions. For each sample, a total of 0.15 µg peptide mixture was subjected to LC-MS analysis. A variety of mass spectrometers were used, including an Orbitrap Lumos Mass Spectrometer or a QE-HF Mass Spectrometer (see above), and a Orbitrap Eclipse and a QE-plus mass spectrometer. All were equipped with a nano-electrospray ion source. Mass spectrometers were operated in DDA mode with an approximate total cycle time of approximately 1 s. Each MS1 scan was followed by high-collision-dissociation (HCD) of the 20 most abundant precursor ions with dynamic exclusion set to 30 seconds. For MS1, 3e6 ions were accumulated in the Orbitrap over a maximum time of 50 ms (25 ms QEplus) and scanned at a resolution of 120,000 FWHM (70,000 FWHM QEplus) at 200 m/z. MS2 scans were acquired at a target setting of 1e5 ions, maximum accumulation time of 50 ms (110 ms QEplus) and a resolution of 30,000 FWHM (17,500 FWHM QEplus) at 200 m/z. Singly charged ions, ions with charge state ≥ 6 and ions with unassigned charge state were excluded from triggering MS2 events. The normalized collision energy was set to 27%, the mass isolation window was set to 1.4 m/z and one microscan was acquired for each spectrum.

#### Protein Identification and Quantification

For affinity pulldown analysis, the acquired raw-files were searched by FragPipe using MSFragger (Kong *et al*, 2017, 201) against *A. baylyi* ADP1 (downloaded from Uniprot 20220109) and 392 commonly observed contaminants. Default settings were used for search, search criteria were set as follows: full tryptic specificity was required (cleavage after lysine or arginine residues unless followed by proline), 2 missed cleavages were allowed, carbamidomethylation (C) was set as fixed modification and oxidation (M) and N-terminal acetylation, as well as phosphorylation (STY) when applicable as a variable modification. A target-decoy search strategy was used to obtain a protein false discovery rate of 1%. Search results were analyzed using the Scaffold software environment (https://www.proteomesoftware.com). Quantitative results were statistically evaluated using the MSstats R package v.4.9.9 (Choi *et al*, 2014).

### Flow-induced dispersion analysis (FIDA)

10 μM TagF was labelled in 20 mM Hepes, 150 mM NaCl, pH 7.4 by reaction for 60 minutes at room temperature with 50 μM DyLight 488 NHS-Ester (Thermo Fisher) prepared as a 1 mM stock in DMSO and diluted into the buffer. Non-conjugated dye was removed by dialysis overnight at 4°C with a Slide-A-Lyzer Dialysis Cassette, 10KDa MWCO (Thermo Fisher) against the same buffer. Flow induced Taylor dispersion experiments were conducted on a Fida Neo (Fidabio) with a 480 nm detector. Measurements were made at 25°C in 20 mM Hepes, 150 mM NaCl, 0.05% Tween 20, pH 7.4, using a coated Fida capillary. For the TagF-Hcp interactions, the analyte solution was a serial dilution of Hcp using a 1:1 mixing ratio, over 12 points starting at 476 μM; and the analyte-indicator solutions were DyLight 488 labelled TagF wt or mutants diluted to 80 nM using the analyte solution from each titration point. **For the TagF-TagZ_1-26_ interaction** (peptide information see TagZ peptide design), the analyte solution was a serial dilution of TagF using a 1:1 mixing ratio, over 12 points starting at 312 μM; and the analyte-indicator solution was FITC labelled TagZ_1-26_ diluted to 100 nM using the analyte solution from each titration point. The capillary was filled with analyte solution at 3500 mbar for 20 sec, followed by the analyte-indicator solution at 50 mbar for 10 sec and measurement at 400 mbar for 180 sec. All data points were measured in triplicate. Taylorgrams were fitted with the standard analysis parameters using the 2 species fit to account for the presence of a small amount of free dye and viscosity compensated using the residence time of the indicator in absence of analyte. The dissociation constant was fitted with a 1:1 model based on the hydrodynamic radius.

### Refeyn Mass Photometry

Mass photometry experiments were performed using a Refeyn OneMP instrument at room temperature (approximately 25 °C). Each measurement was made by mixing 2 µl of sample into 18 µl of buffer in a droplet contained by a 6-well silicone sample cassette (Refeyn) on 24×50 mm area 170±5 µm thickness coverglass (Marienfeld). Data acquisition was performed using the instrument control software AcquireMP 2024 R2. The stock solution of the MassFerence P1 protein standards (Refeyn) was diluted 500-fold in 20 mM phosphate, 150 mM NaCl, pH 7.5 (PBS) and interferometric scattering contrast movies were recorded for 120 s for instrument calibration. 5 samples of Hcp were diluted to a final concentration of 38 nM in PBS and interferometric scattering contrast movies were recorded for 120 s.

The resulting movies were analysed to obtain contrast event histograms in the analysis software DiscoverMP 2024 R2 with the default analysis settings. Calibration curves relating contrast to molecular mass were constructed using the data from the MassFerence P1 protein standards and the automated routine in the software. For each sample of Hcp, the most recent calibration curve was applied to the contrast event histogram to give a molecular mass event histogram, which was plotted using a bin-width of 5 kDa. To obtain a mean and standard deviation for the molecular mass of each discrete species in the sample, each peak in the molecular mass event histogram was fitted to Gaussian using the automated peak-fitting procedure in the software.

### Negative Stain Electron Microscopy

Diluted sample (1:50, 5 µL) was adsorbed for 60 s onto glow-discharged (30 s / 20 mA, Quorum GloCube, UK) 400-mesh copper grids (G-400Cu, Electron Microscopy Sciences) coated with ∼4 nm parlodion/carbon (Safematic, CH). The grids were stained with 2% uranyl acetate, and subsequently imaged with a Talos L120C electron microscope (Thermo Fisher Scientific, USA) operating at 120 kV (92k x magnification, 1.54 Å/pix), equipped with a CETA CMOS camera.

### TagF-TagZ interaction assays

#### TagZ peptide design

Peptide sequence was initially based on TOPCONS (Tsirigos *et al*, 2015) trans-membrane region prediction, but 5 additional C-terminal residues were included in the sequence to ensure the complete motif would be present on the sequence. A glycine-serine linkage was added to the peptide C-terminus to increase distance the fluorescein label to the putative recognition site on the peptide. The hydrophobicity score was calculated using Expasy ProtScale (Gasteiger *et al*, 2005) with the Kyte and Doolite model and a sliding window of 7 residues. Peptides were ordered with and without C-terminal fluorescein label but otherwise identical sequence from peptides & elephants GmbH:

| peptide | sequence | Nt/Ct | counter-ion | MW calc g/mol |
| --- | --- | --- | --- | --- |
| unlabeled | H-<br>MAQEKPTSLRILQTPKKKSVSPLLFS<br>GS-OH | H/OH | HCl | 3073.0 |
| fluorescein-labelled | H-<br>MAQEKPTSLRILQTPKKKSVSPLLFS<br>GS (FITC)-OH | H/OH | HCl | 3590.3 |

#### Binder design

Binders were designed against the crystal structure of TagF. We used RFdiffusion (Watson *et al*, 2023) (model BFF_9) with 50 denoising time steps for 75-residue-long minibinder backbone structure design, specifying hotspots W36, E234 and D255 on TagF to condition for design of binders interacting with the desired target site. Backbones generated by RFdiffusion were sequence designed using ProteinMPNN (Dauparas *et al*, 2022) (t=0.1), generating 10 sequences per backbone.

We used AlphaFold2 (Jumper *et al*, 2021) and initial guess (Bennett *et al*, 2023) to predict structures from the designed sequences. The resulting structural library was filtered based on AlphaFold2 metrics pAE interaction and AlphaFold2 pLDDT_binder (minibinder in complex with target) and metrics calculated by Rosetta (Leaver-Fay *et al*, 2011): Cα RMSD between binder in complex with target vs binder alone; spatial aggregation propensity (SAP) score; and change in free energy upon binding (ddG).

To improve design metrics, partial diffusion (Vázquez Torres *et al*, 2024) was used to further optimize structure and sequence of the best scoring designs.

This *in silico* design-test-optimization cycle was repeated until 24 binder designs passed the following filters:

| Set for experimental testing | pLDDT binder | pLDDT binder_mono | pAE interaction / Å | C<sub>α</sub> RMSD / Å | SAP | ddG / kcal/mol |
| --- | --- | --- | --- | --- | --- | --- |
| 24 designs | > 94 | > 93 | < 3.8 | < 0.6 | < 44 | < -75.55 |

DNA Sequence used for cloning and expression of binder F2C1:

ATGAGCGCCGAAGAAGAACTGAAAAACGCGAAACAGGGCGCGGAAAACCTGCTGGAAGGCACCAAAAACCGCGTTAAAGAACTGAAAGCGGAAGGCGAAAGCACCAAAACCCTGATTGCGCAGGTGAACATCAAAGCGCTGTATGCGGAATATCTGGCGGAAAAATACGGCTATGAAGTGTATAAAGAAACCGCCAAAAAACTGAAAGAACTGGCGAAAGAACTGGAAGGAAGCGGTGGTTCAGGATGGAGCCACCCGCAGTTCGAAAAGTAA

## Funding

M.K.E., A.H. and F.D. were supported by Biozentrum PhD fellowships. This work was funded by the European Research Council grant no. 865105 and the Schweizerischer Nationalfonds zur Förderung der Wissenschaftlichen Forschung (Swiss National Science Foundation), grant no. 10000935 awarded to M.B..

We thank Dr. Timothy Sharpe from the Biophysics Facility (University of Basel) for guidance with experiment design, data collection and analysis, as well as Dr. Thomas Bock from the Proteomics Facility (University of Basel) for support with data acquisition and analysis. We thank Dr. Roman Jakob for help with X-ray data collection. We also thank Dr. Moritz Hunkeler from the BioEM Lab (University of Basel) for assistance with negative stain electron microscopy.

## Supplementary Figures

**Supplementary Figure 1:**
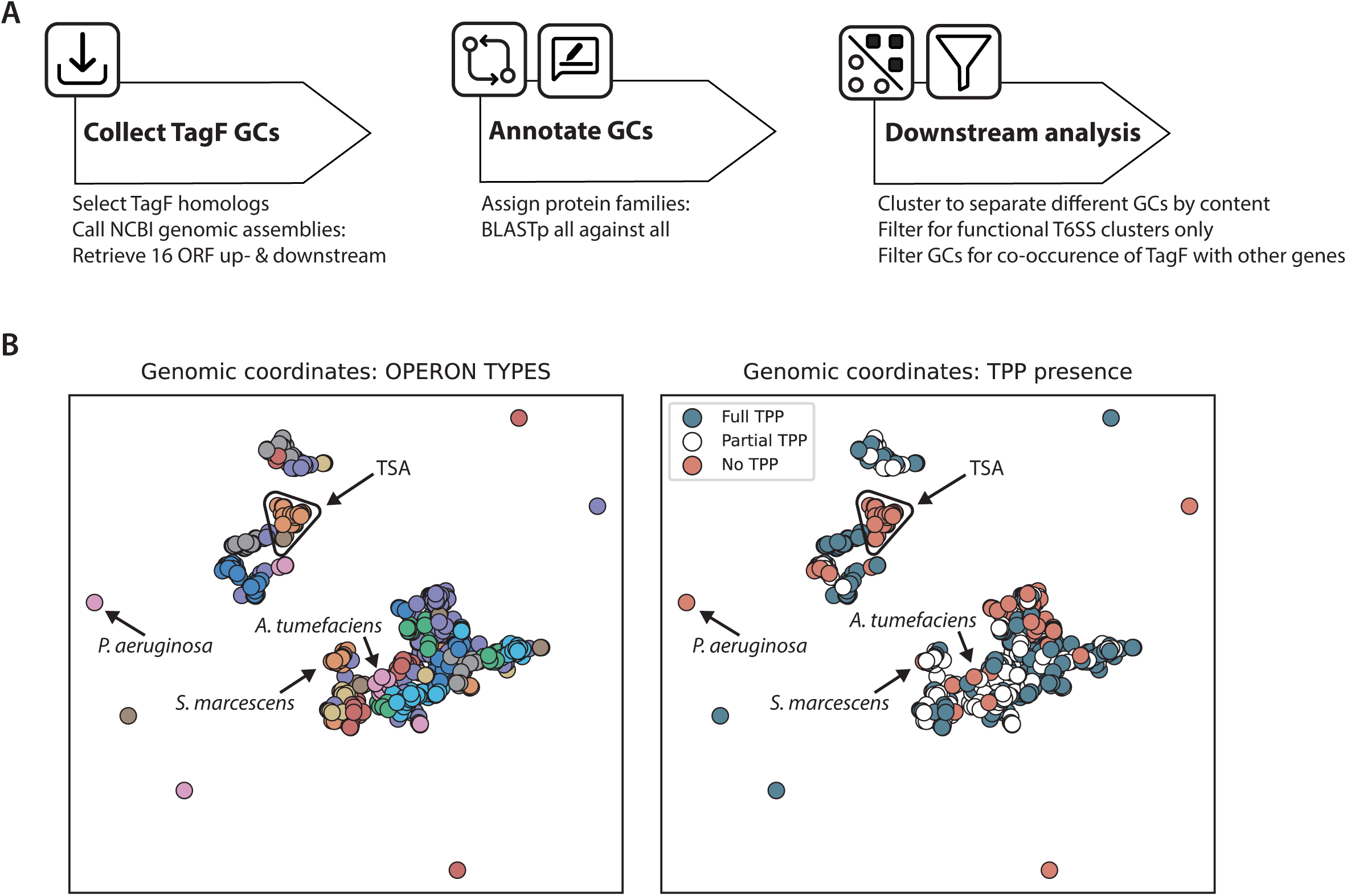
GCSnap analysis of TagF genomic contexts. **A** GCSnap workflow: Based on a diverse list of TagF protein sequences, 16 ORF on either side of the gene are retrieved and consecutively annotated based on sequence homology. Downstream analysis includes filtering based on gene content and clustering approaches as shown in (B). **B** GCSnap genomic context coordinates. GCs clustering according to content, panels show T6SS loci colored according to operon type (left) and TPP presence-absence (right). Box and arrow indicate loci type including *Acinetobacter*, TPP-type *tagF* loci of interest are indicated.

**Supplementary Figure 2:**
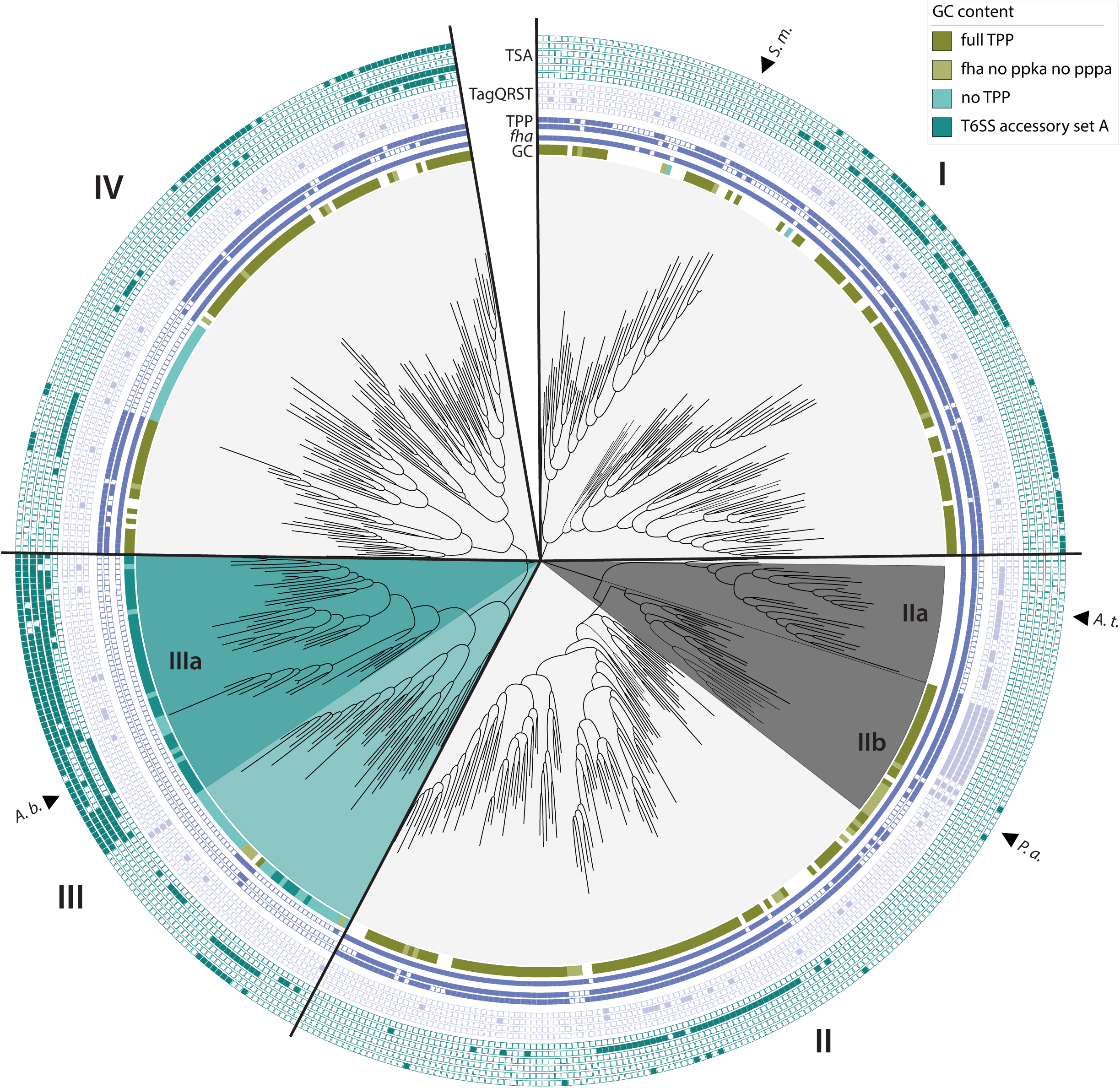
TagF phylogenetic tree. Phylogenetic analysis of TagF sequences from GC analysis visualized using iTOL with annotations based on GCSnap analysis. From the inside out: Categorization according to TPP/TSA (legend top right); Purple: Fha, PpkA, PppA; Light purple: TagS, TagT, TagQ, TagR; Teal: TsmK, TsmK-like, TagX, TagZ, TslA, TagN. The fully annotated tree exported at high resolution is available in Supplementary File 2. TsmK and TsmK-like homologs (lacking a transmembrane region) were combined for the frequency plots in Figure 1A.

**Supplementary Figure 3:**
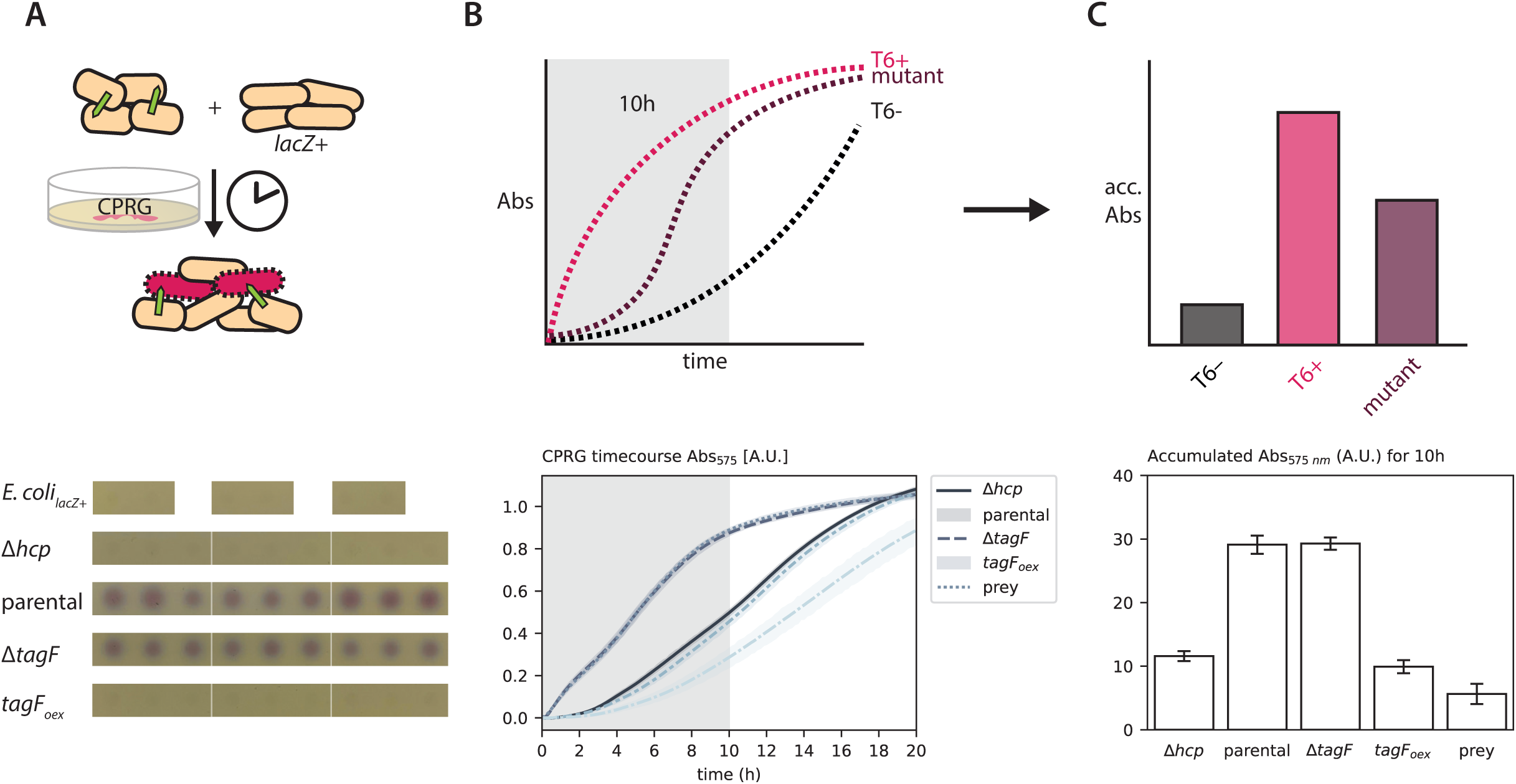
CPRG accumulation assay. **A** Top: Cartoon of T6SS CPRG lysis assay workflow. *A. baylyi* T6SS+ strains or T6SS mutant strains are mixed with susceptible *lacZ+ E. coli and* plated on CPRG LB-agar. Bottom: Example showing killing mixes prepared in biological and technical triplicate and spotted agar using a pin tool. **B** Lysis assay 20 h time course monitoring chlorophenol-red indicator absorbance at 575 nm of a plate corresponding to (A). Impaired mutant phenotypes result in intermediate lysis activity. **C** Accumulative lysis from absorbance reads for the first 10 h of the experiment. The corresponding area of the time course plot is indicated in light grey in (B).

**Supplementary Figure 4:**
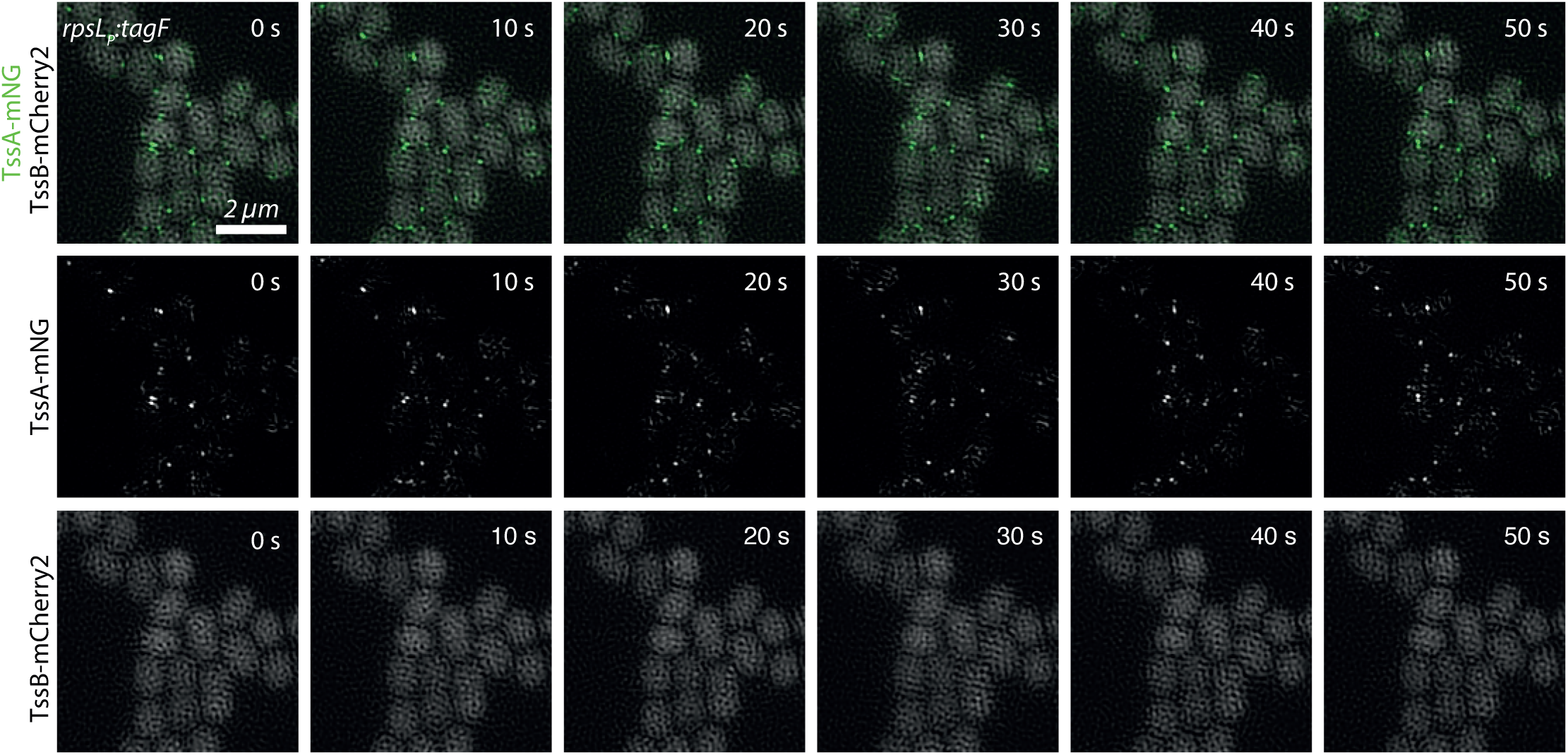
TssA-mNG labelling. 2-color time-lapse 2D-SIM imaging *TagF_oex_*sheath (TssB-mCherry2) and TssA-mNG localization as in Figure 1G, showing single channel panels. TssB-mCherry2 signal-to-noise was not sufficient to reconstruct contact site foci observed for TssB-msfGFP.

**Supplementary Figure 5:**
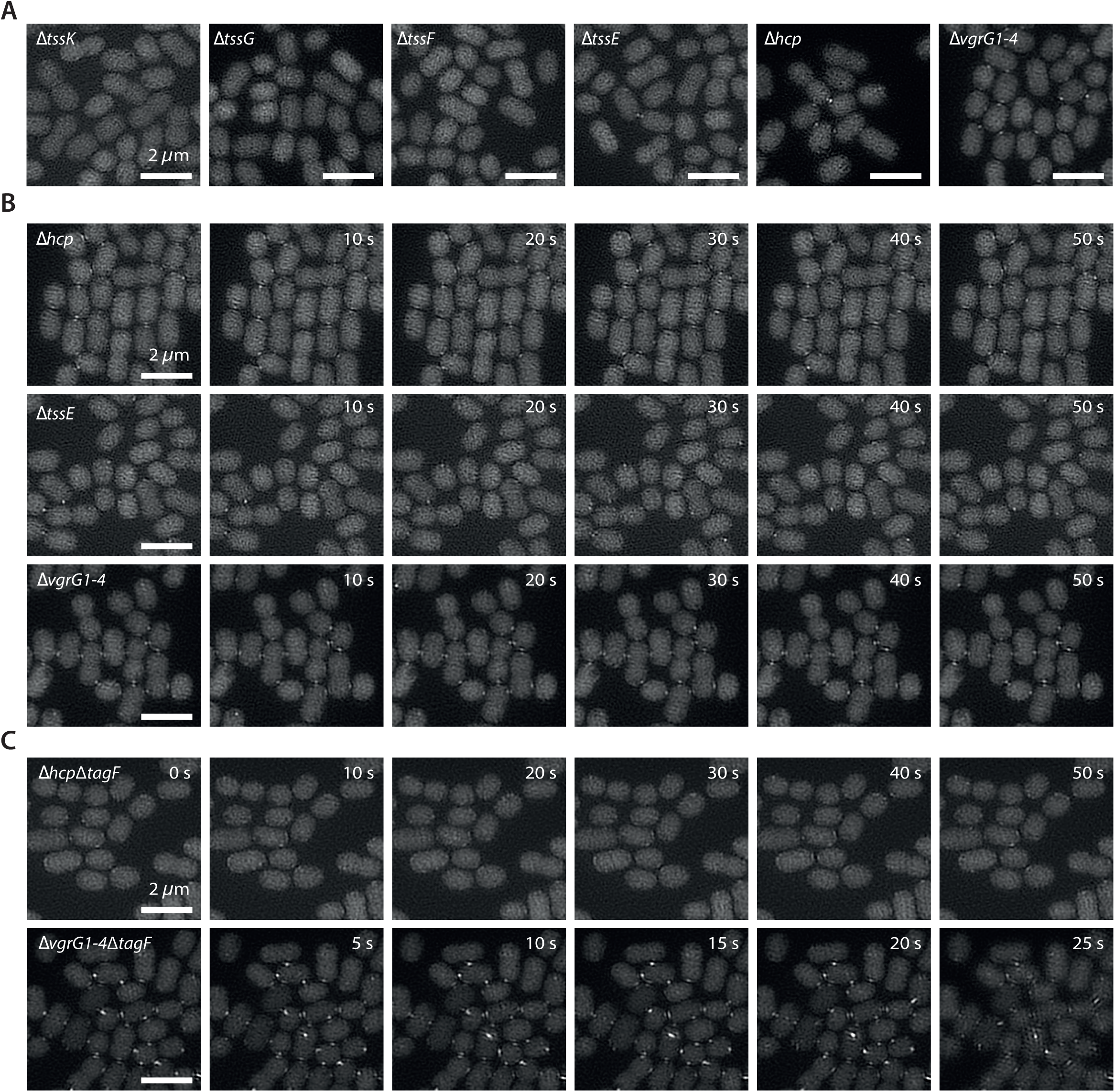
SIM of baseplate mutants to identify TagF regulatory target. **A** 2D-SIM of sheath localization (TssB-sfGFP) in the different baseplate mutants screening for sheath foci. **B** Time-lapse 2D-SIM following sheath dynamics (TssB-sfGFP) of *Δhcp, ΔtssE, and ΔvgrG1-4* mutant strains. **C** Time-lapse 2D-SIM following sheath dynamics (TssB-sfGFP) of *tagF* deletion strains in *Δhcp* and *ΔvgrG1-4* backgrounds.

**Supplementary Figure 6:**
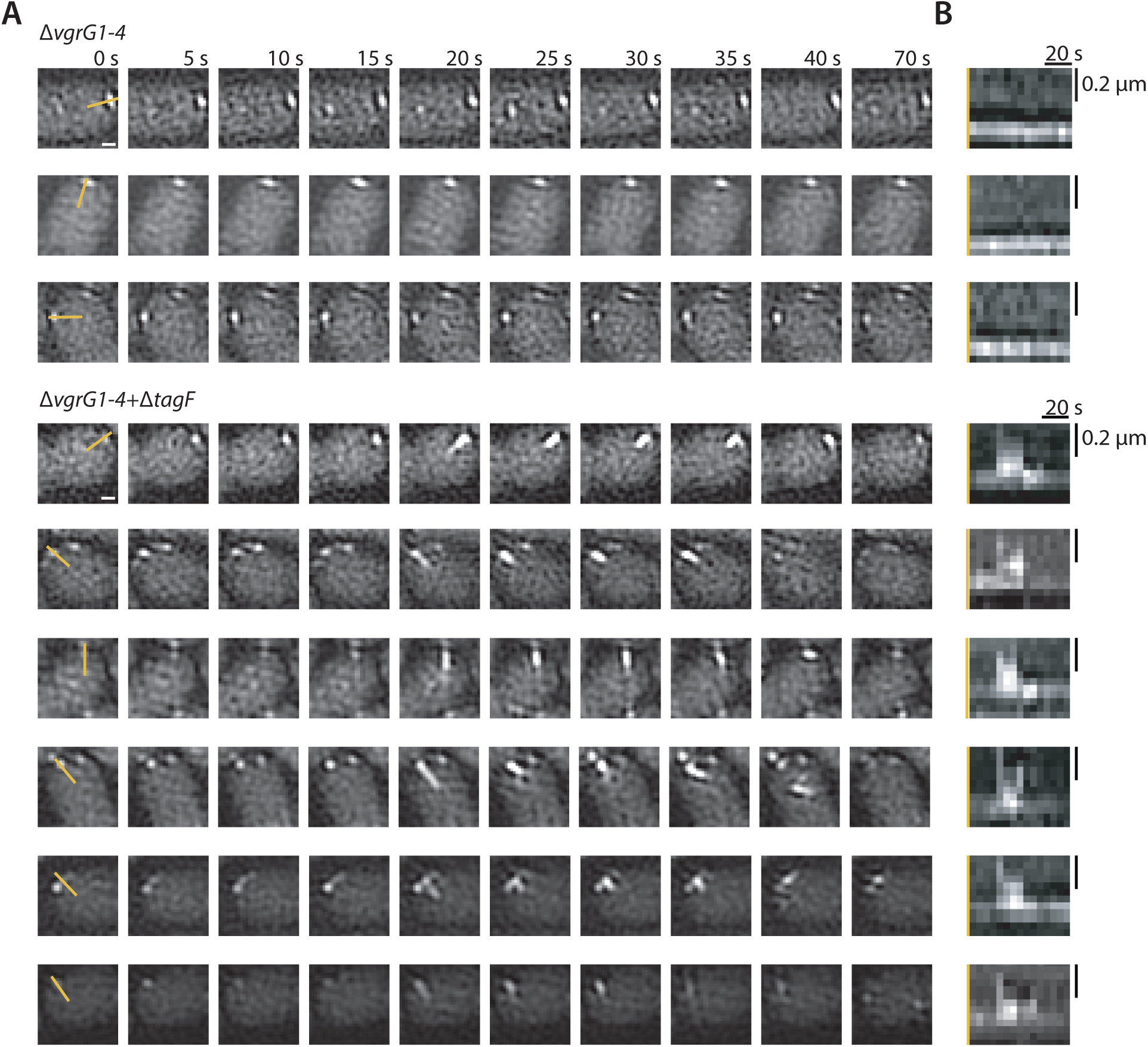
Additional examples for precursor mutant sheath dynamics: SIM timelapses and corresponding kymographs. **A** Additional examples of 2D-SIM time-lapse following *ΔvgrG1-4* and *ΔvgrG1-4*Δ*tagF* sheath dynamics (TssB-sfGFP). Scale bar represents 0.2 µm. Yellow lines in the first panels indicate kymograph origin in (B). **B** Kymographs of the time-lapse imaging in (A).

**Supplementary Figure 7:**
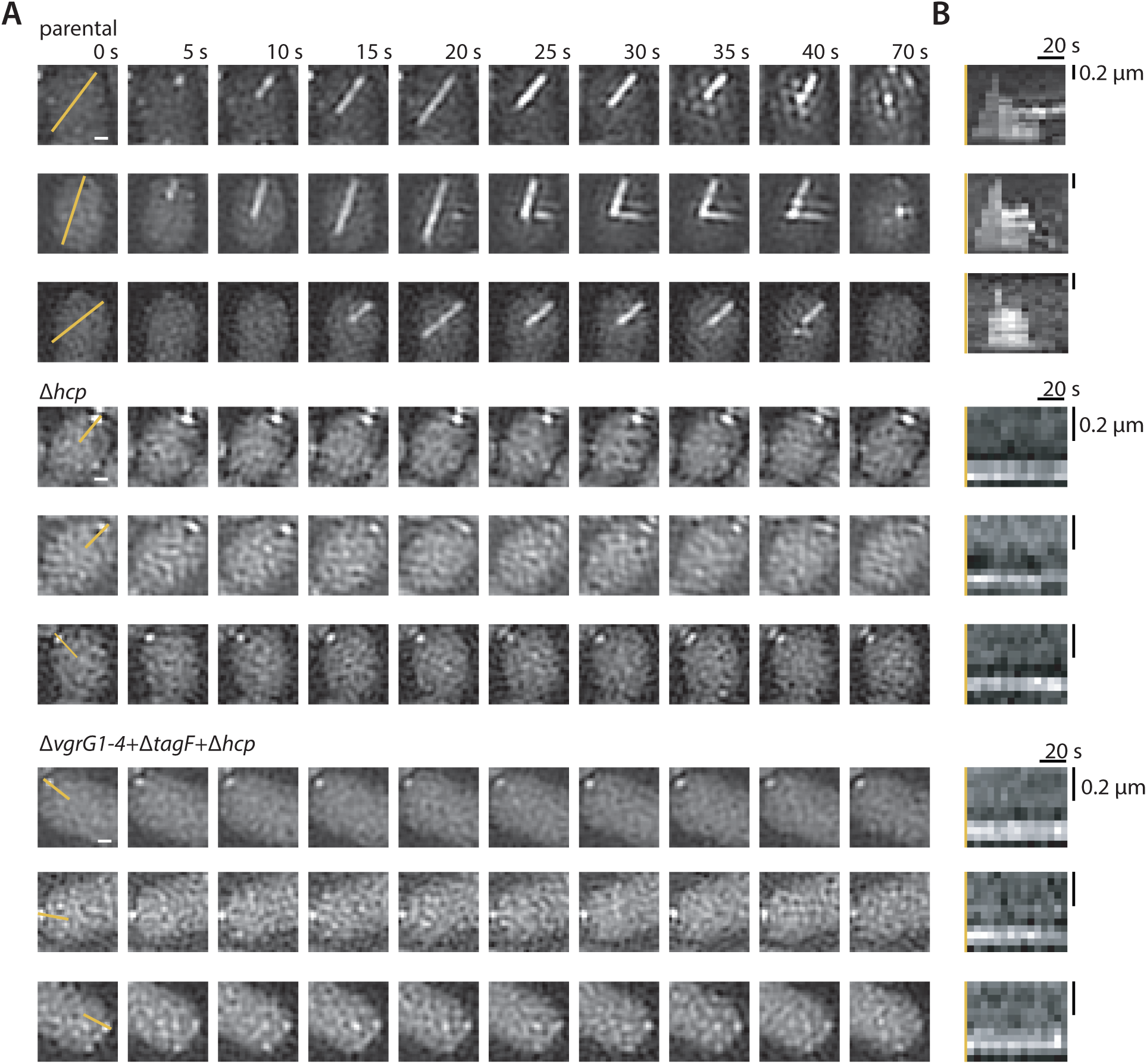
Additional examples for parental strain and *Δhcp* sheath dynamics: SIM timelapses and corresponding kymographs. **A** Additional examples of 2D-SIM time-lapse following parental strain and *Δhcp* mutant sheath dynamics (TssB-sfGFP). Scale bar represents 0.2 µm. Yellow lines in the first panels indicate kymograph origin in (B). **B** Kymographs of the time-lapse imaging in (A).

**Supplementary Figure 8:**
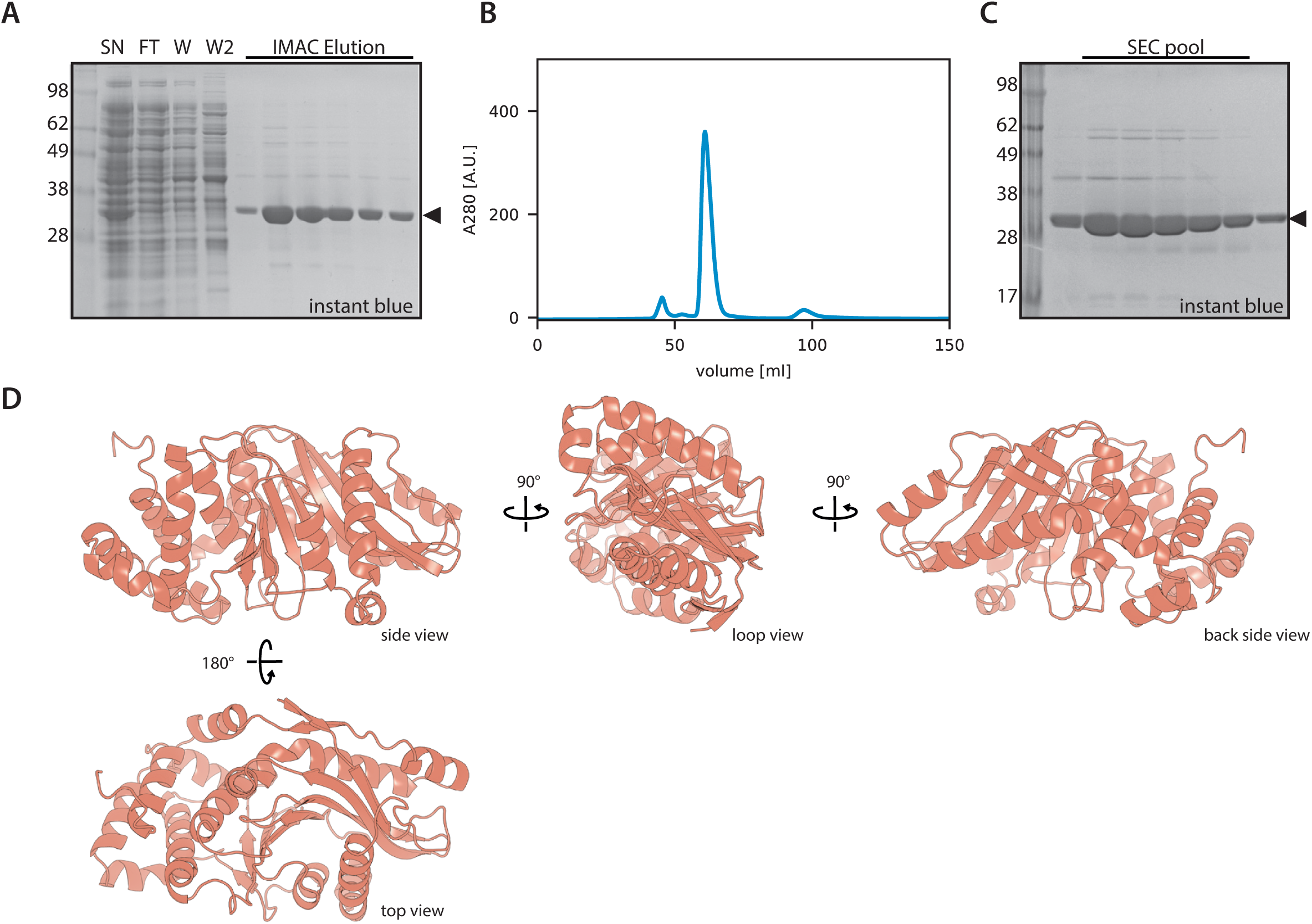
TagF purification and alternative views of the TagF structure. **A** SDS-PAGE of TagF-His_10_ purification stained with instant blue. SN: ultracentrifugation supernatant, IMAC-purification: FT flowthrough, W1: washes with purification buffer W2: wash with 50 mM imidazole, E: elution using 300 mM imidazole. **B** SEC chromatogram of TagF after His_10_-tag cleavage. Column: Superdex 75 HiLoad 16/600 pg. **C** SDS-PAGE of TagF SEC elution fractions. **D** Different views of *A. baylyi* crystal structure in cartoon representation.

**Supplementary Figure 9:**
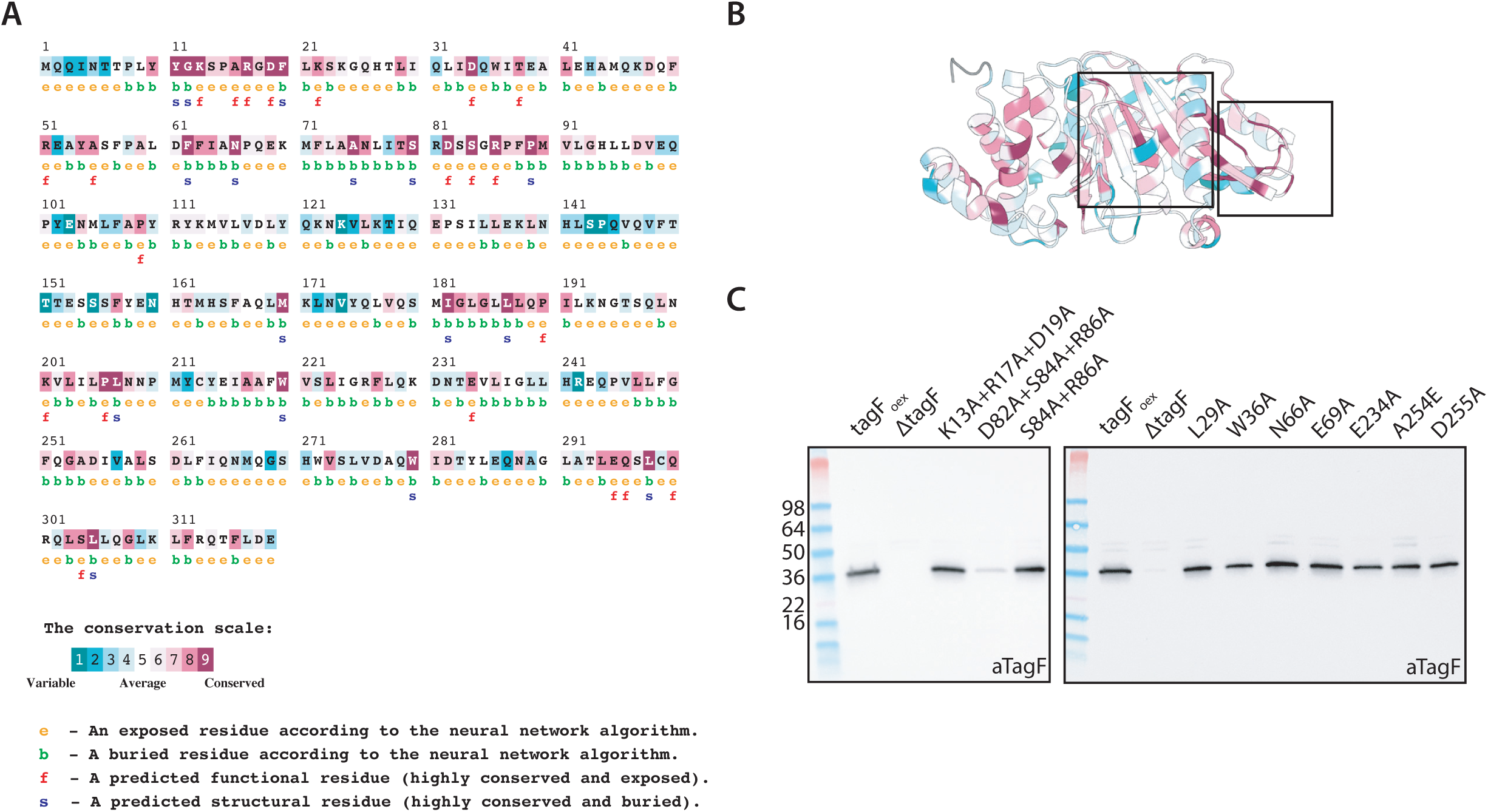
TagF sequence conservation and point mutant expression checks. **A** ConSurf sequence analysis of TagF based on TSA-type TagF sequences. **B** TagF cartoon representation colored using sequence conservation color scheme from ConSurf analysis (A). **C** Full western blots (Figure3D) from protein expression analysis of TagF mutants from the *A. baylyi* soluble lysis fraction using specific anti-TagF polyclonal antibody.

**Supplementary Figure 10:**
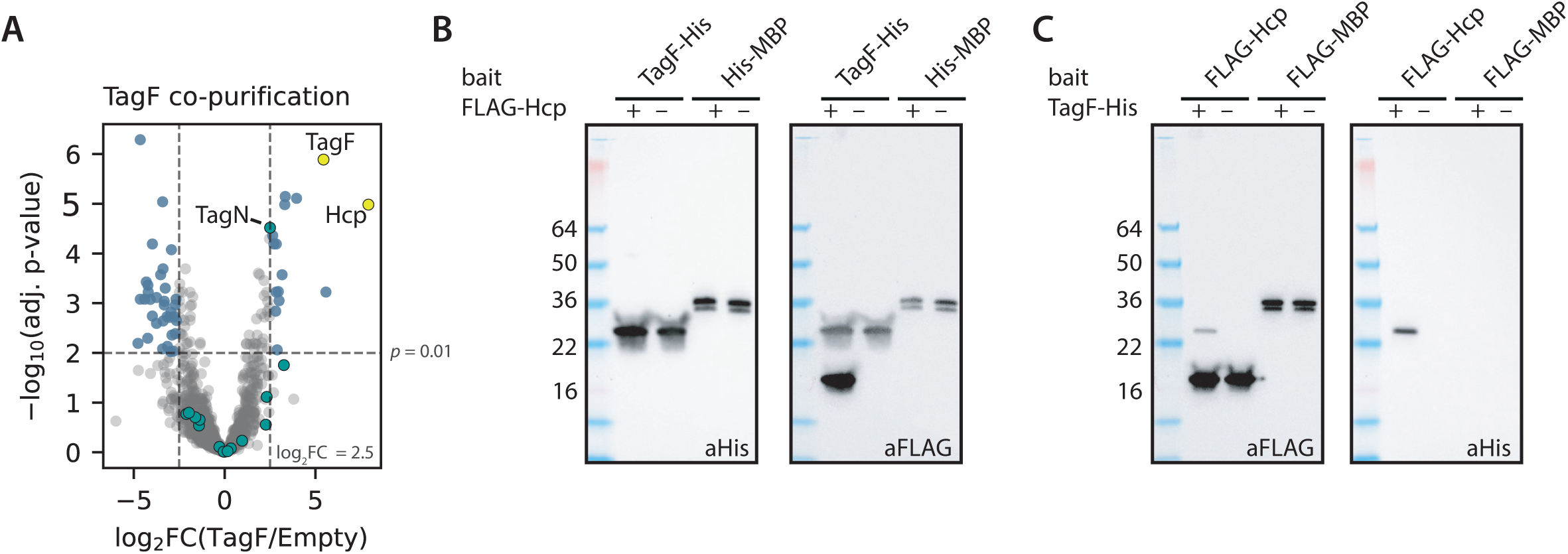
TagF interaction screen identifies Hcp. **A** LC-MS/MS statistical analysis of TagF co-purification. The volcano plot shows log_2_-fold change of protein enrichment using TagF-His_10_ bait versus empty beads. Cutoffs shown are log_2_FC = 2.5 and *adj. p* = 0.01. Hcp is significantly enriched in the sample with TagF bait. Yellow: TagF and Hcp; Teal: other T6SS proteins; Blue: non-T6SS proteins. **B** Full western blot of co-purification from *E. coli* using TagF-His bait and FLAG-Hcp prey. His-MBP was used as control-bait. **C** Western blot of co-purification from *E. coli* using FLAG-Hcp bait and TagF-His prey. FLAG-MBP was used as control-bait.

**Supplementary Figure 11:**
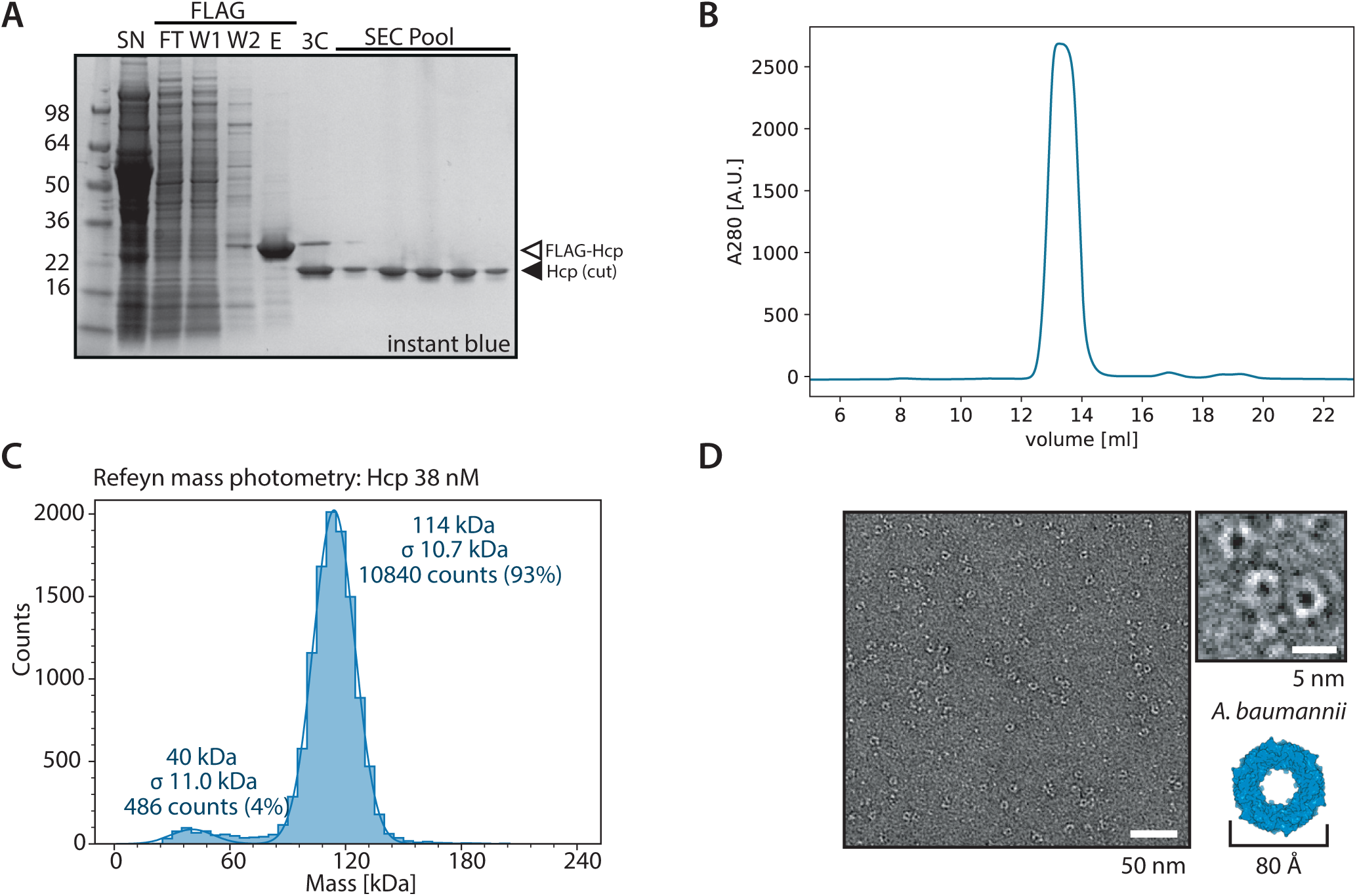
Purification and characterization of *A. baylyi* Hcp. **A** SDS-PAGE of Hcp-FLAG purification stained with instant blue. SN: ultracentrifugation supernatant; FLAG-purification: FT: flowthrough; W1, W2 : washes with purification buffer; E: elution pool. **B** Hcp SEC purification chromatogram; column: Superdex 200 increase 10/300 gl. **C** Refeyn mass photometry analysis of pure Hcp sample (5 replicates) indicates that at least 93% of Hcp is present as hexameric complex. **D** Negative stain electron micrograph of Hcp sample. Particle shape corresponds to the expected hexameric ring structure of the free Hcp building blocks of T6SS tube. Inset shows the *A. baumannii* Hcp hexameric structure for comparison (PDB 4w64).

**Supplementary Figure 12:**
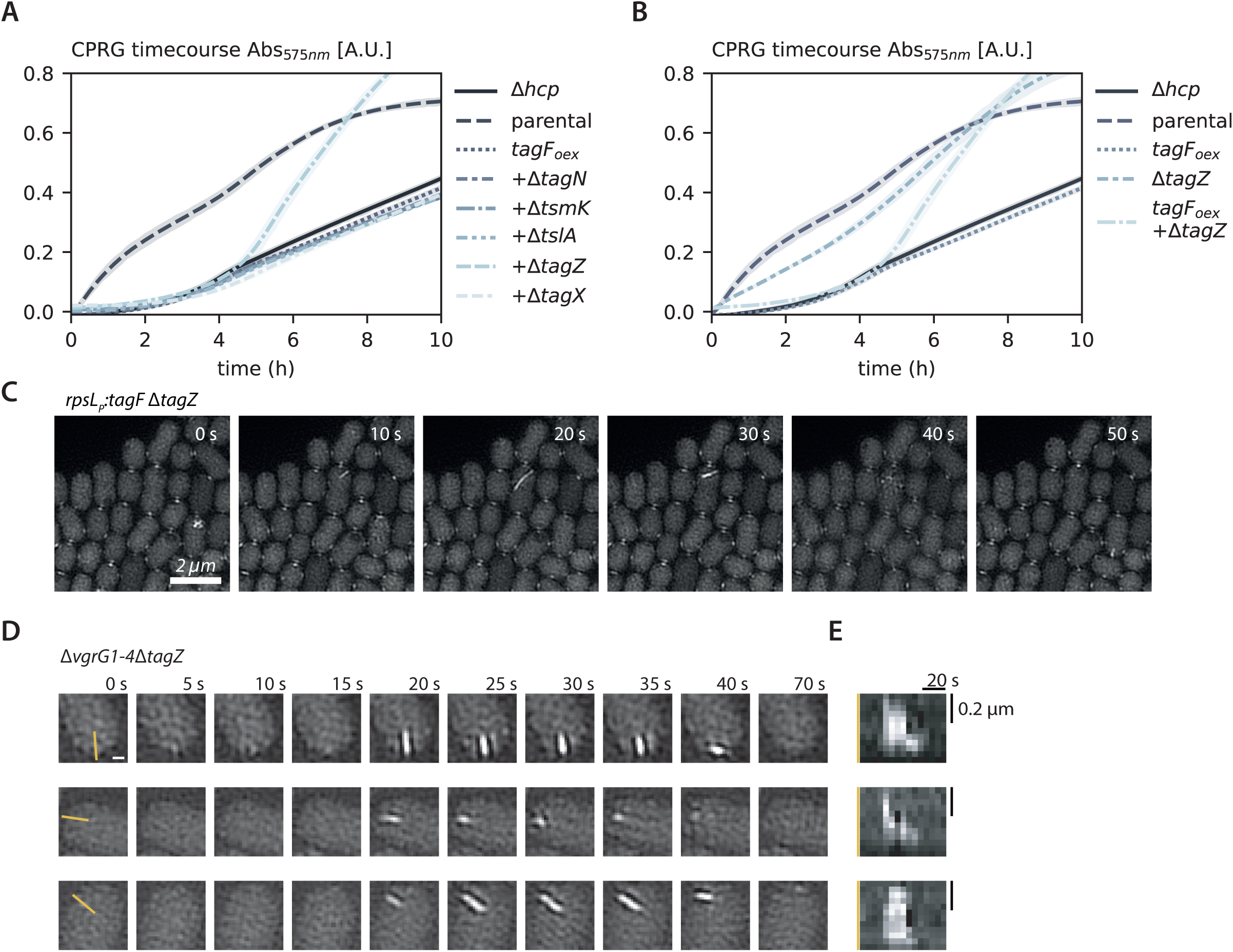
TagZ is required to maintain TagF-mediated function. **A** CPRG lysis assay time course comparing T6SS-dependent *E. coli* lysis of strains overexpressing TagF in accessory gene deletion backgrounds. **B** CPRG lysis assay time course comparing T6SS-dependent *E. coli* lysis of strains TagF_oex_ *± TagZ* and *ΔtagZ*. **C** Time-lapse 2D-SIM following sheath dynamics (TssB-sfGFP) of *TagF_oex_* in absence of *tagZ*. **D** Additional examples of 2D-SIM time-lapse following *ΔvgrG1-4* and *ΔvgrG1-4*Δ*tagF* sheath dynamics (TssB-sfGFP). Scale bar represents 0.2 µm. Yellow lines in the first panels indicate kymograph origin in (E). **E** Kymographs of the time-lapse imaging in (D).

**Supplementary Figure 13:**
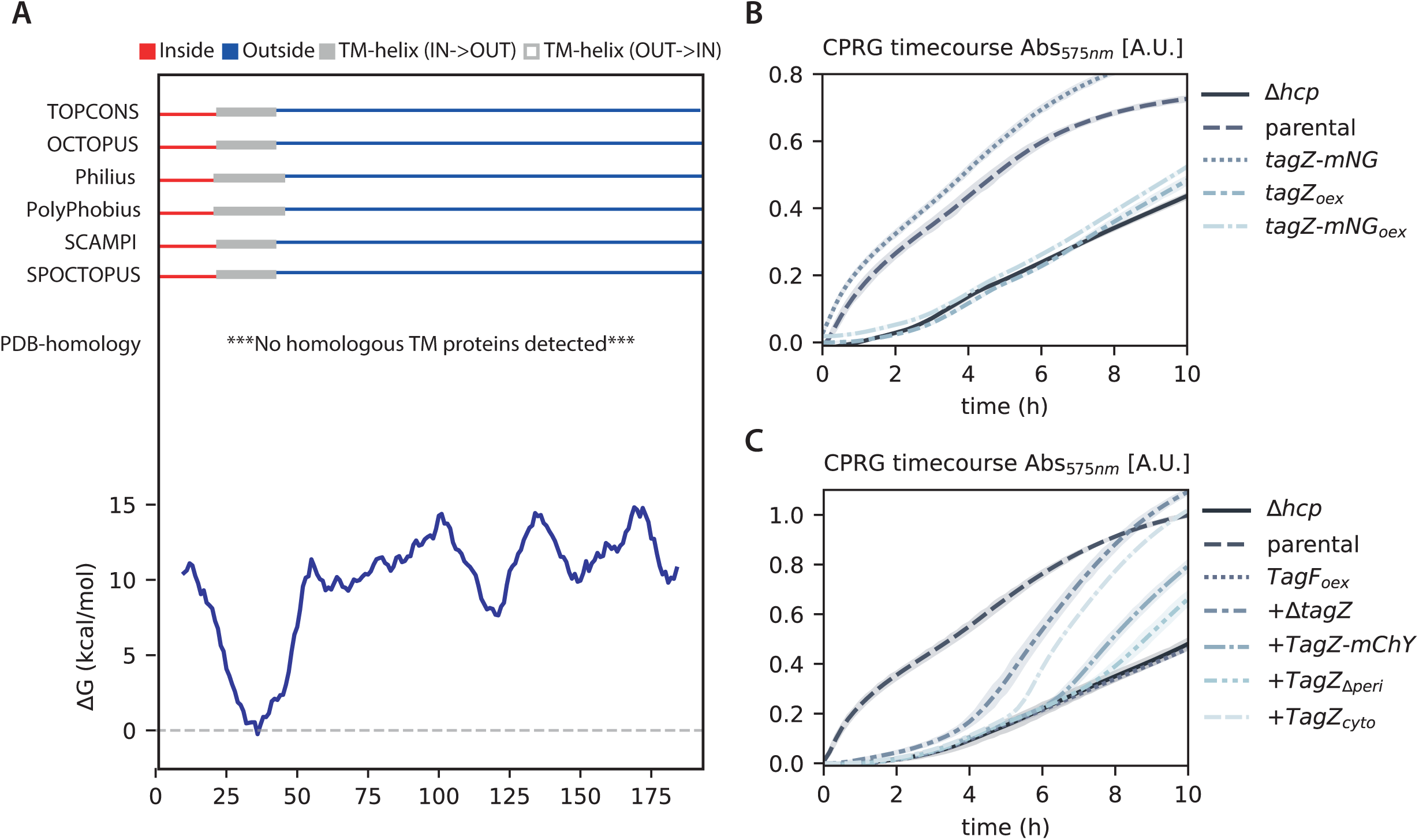
TagZ predicted topology and TagZ construct complementation. A Topcons output for TagZ topology prediction. **B** CPRG lysis assay time course comparing T6SS-dependent *E. coli* lysis of TagZ-mNG strains. **C** CPRG lysis assay time course comparing T6SS-dependent *E. coli* lysis of TagF_oex_ phenotype in strains expressing TagZ truncation constructs.

**Supplementary Figure 14:**
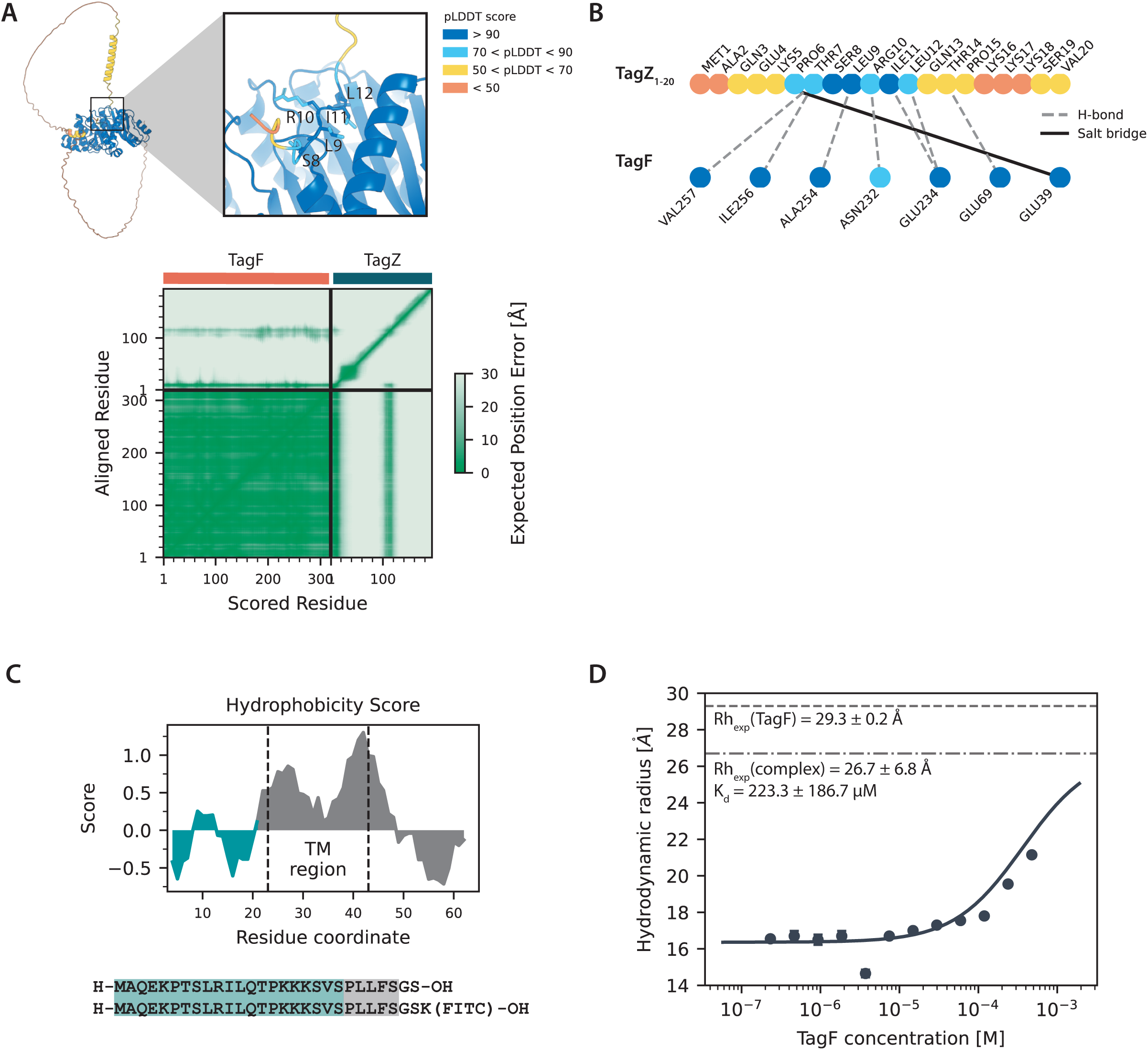
TagF-TagZ AlphaFold model. **A** AlphaFold3 model of TagF-TagZ complex shown in cartoon representation and colored by pLDDT score. Below: AlphaFold3 TagF-TagZ model PAE plot. **B** Labelled interaction network based on PDBePISA analysis, colored based on pLDDT score. **C** TagZ_1-26_ N-terminal peptide design for binding tests. Peptide length was chosen based on TagZ hydrophobicity score to ensure peptide solubility in aqueous buffer. Peptide was designed with C-terminal flexible glycine-serine linker and optional fluoresceine (FITC) modification for binding assays. **D** FIDA analysis of TagZ_1-26_ (FITC) peptide (100 nM) interaction with TagF following complex formation by measuring Rh shows weak binding. Experimental Rh for pure TagF is indicated with a dashed line.

**Supplementary Figure 15:**
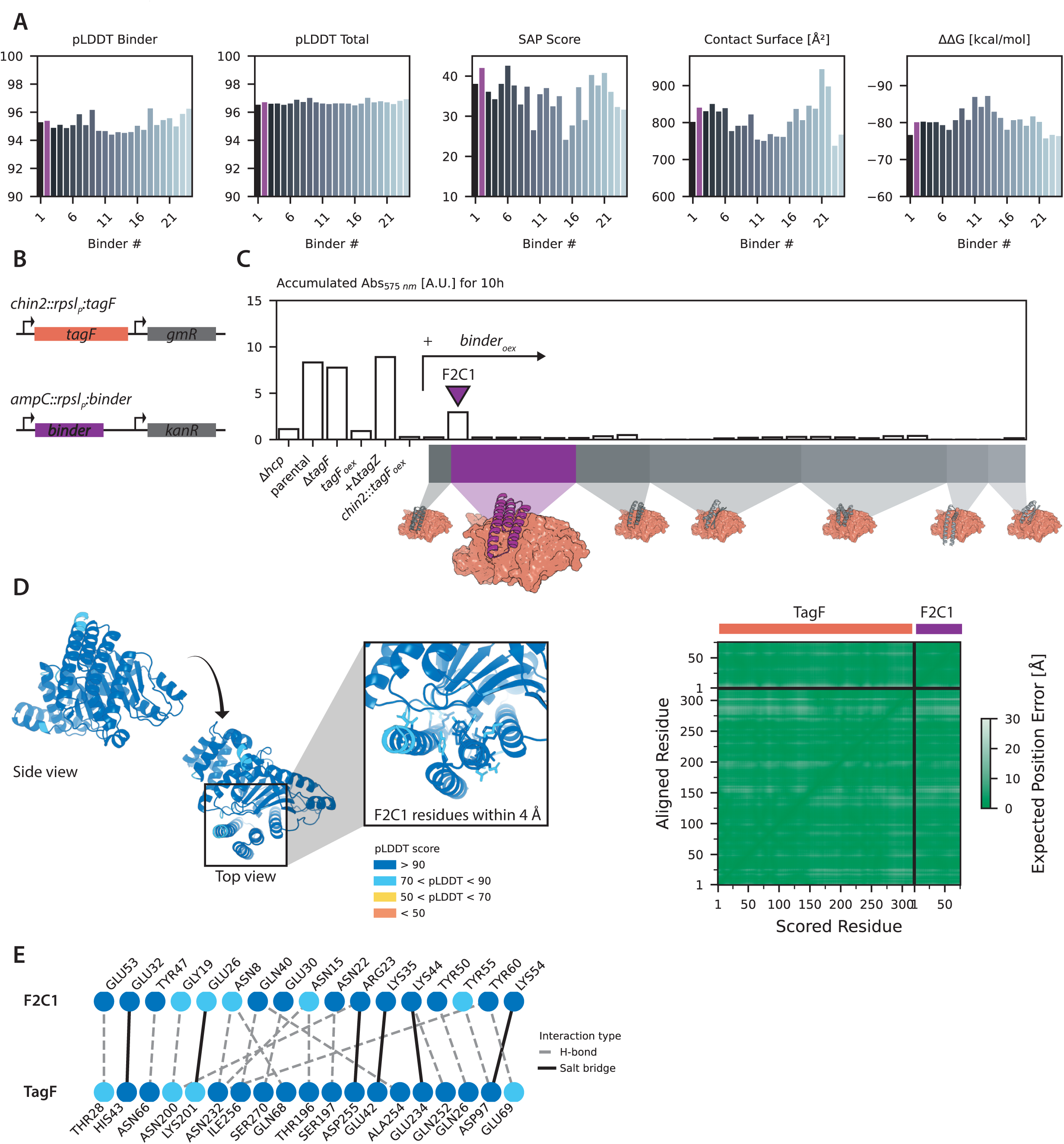
Binder design and screening, F2C1 Alphafold3 confidence. **A** Key statistics for TagF-groove binders selected for phenotypic screen in *A. baylyi*. F2C1 is highlighted in purple. **B** Genomic insertion cassettes designed for co-overexpression of TagF from *chin2* and binders from the neutral *ampC* integration site. The cassettes use the same promoter but distinct antibiotic selection markers. **C** TagF groove-binder functional screen based on CPRG accumulation assay, to identify binders interfering with the T6SS-negative TagF_oex_ phenotype. The different back bone folds of the tested binders are shown in cartoon representation at the TagF-groove interface (orange, surface representation). **D** AlphaFold3 model of TagF-F2C1 binder complex shown in cartoon representation and colored by pLDDT score. Right panel: AlphaFold3 TagF-F2C1 model PAE plot. **E** Labelled interaction network based on PDBePISA analysis, colored based on pLDDT score.

**Supplementary Figure 16:**
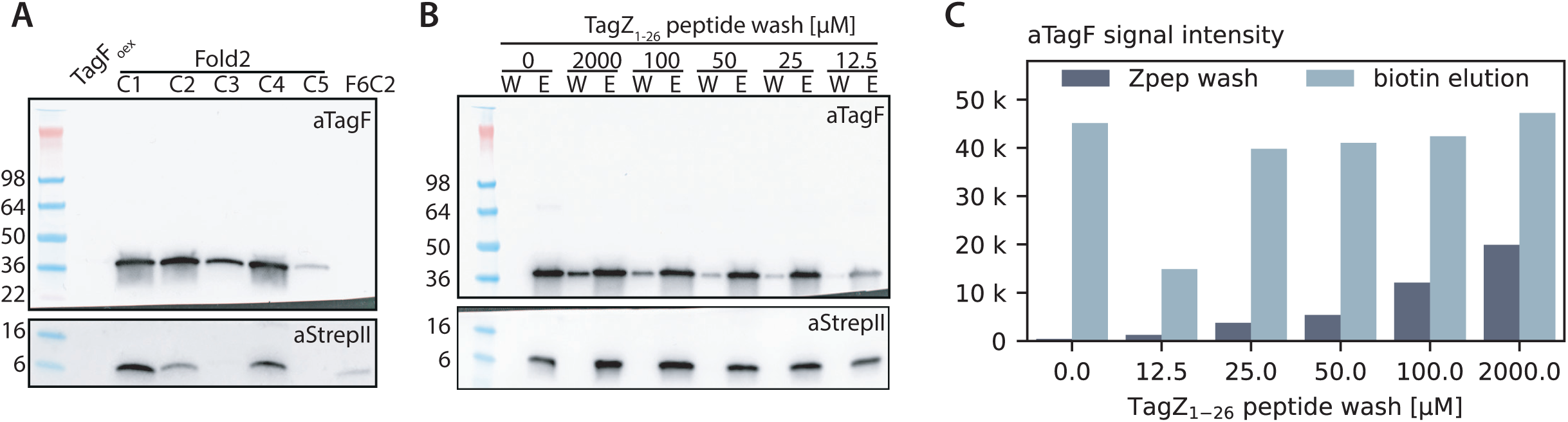
TagF-F2C1 binder co-purification results and the effect of TagZ_1-26_ peptide on the interaction. **A** Full western blot of strepII-tag based co-purification of F2C1-strepII enriched for TagF, as in Figure 5F. **B** Full western blot of strepII-tag based co-purification of F2C1-strepII, challenged with TagZ_1-26_ peptide, as in Figure 5G. W: TagZ_1-26_ wash; E: strepII-tag specific 50 mM biotin elution. **C.** Plot of quantified absolute anti-TagF signal intensities from the blot in (B), basis for the relative signal intensities shown in Figure 5H.

## Supplementary Videos

**Supplementary Video 1: Overexpression of TagF blocks sheath dynamics.**

Time-lapse 2D-SIM following sheath dynamics (TssB-sfGFP) of the parental strain (1-3) and TagF_oex_ (4-6). Scale bar represents 1 µm.

**Supplementary Video 2: TagF controls the spike insertion checkpoint.**

Time-lapse 2D-SIM following sheath dynamics (TssB-sfGFP) of mutants *ΔvgrG1-4 (1-2)*, *ΔvgrG1-4ΔtagF* (3-5) and *ΔvgrG1-4ΔtagFΔhcp* (6-7). Scale bar represents 1 µm.

**Supplementary Video 3: TagZ is required to maintain the spike insertion checkpoint.**

Time-lapse 2D-SIM following sheath dynamics (TssB-sfGFP) of *ΔvgrG1-4ΔtagZ* mutant (1-3). Scale bar represents 1 µm.

## Supplementary Tables related to materials and methods

**Supplementary Table 1:**
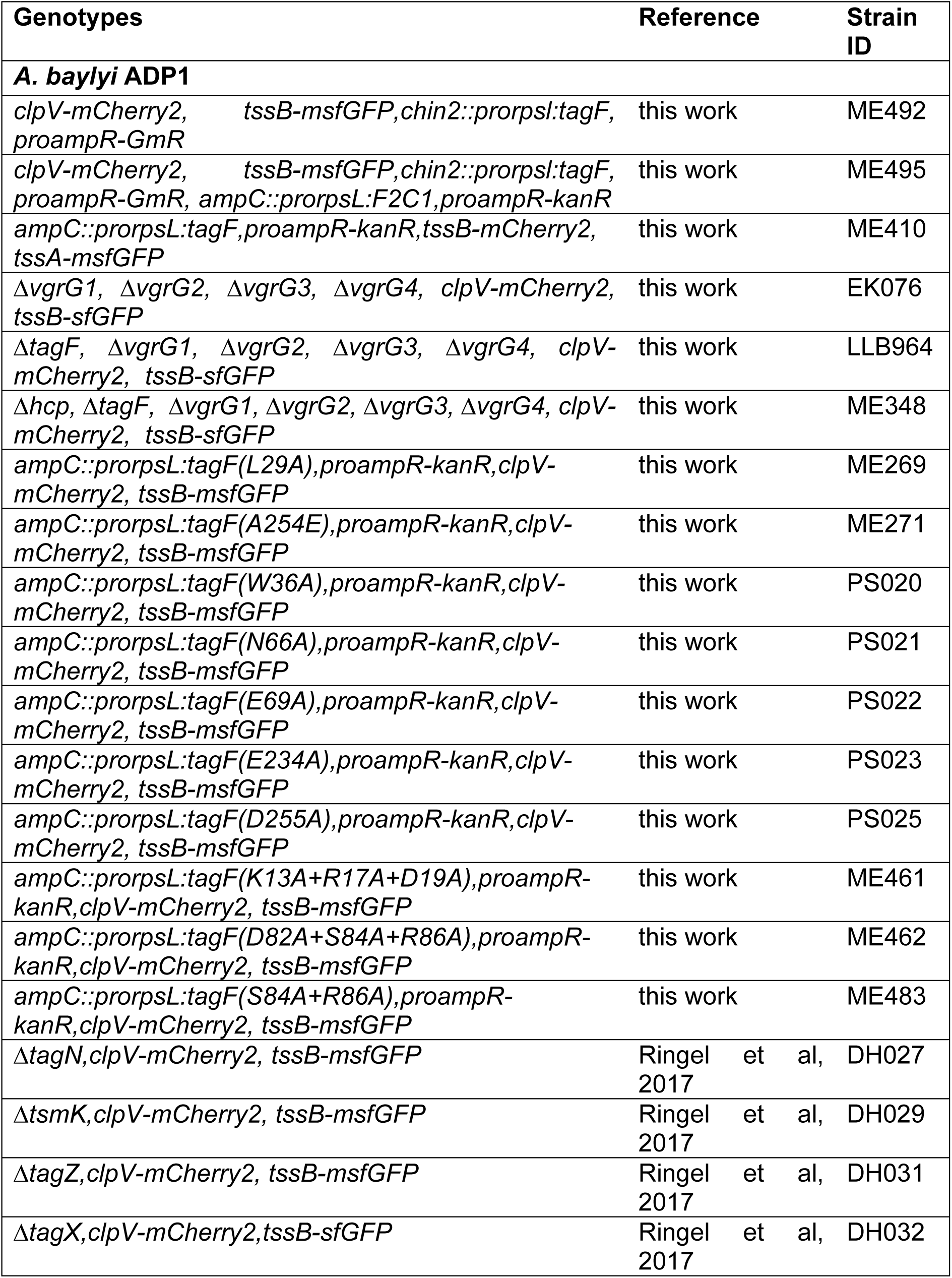
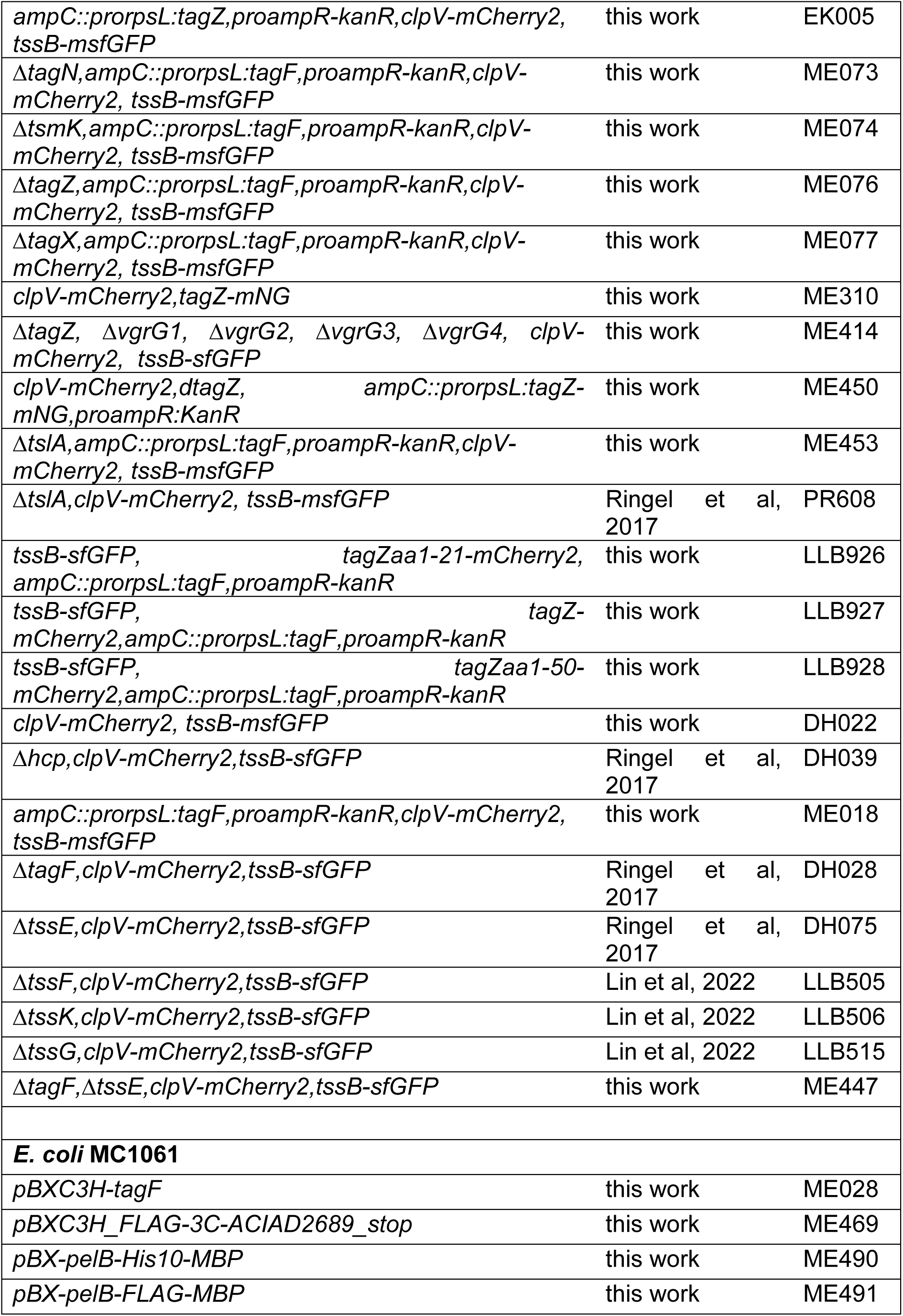
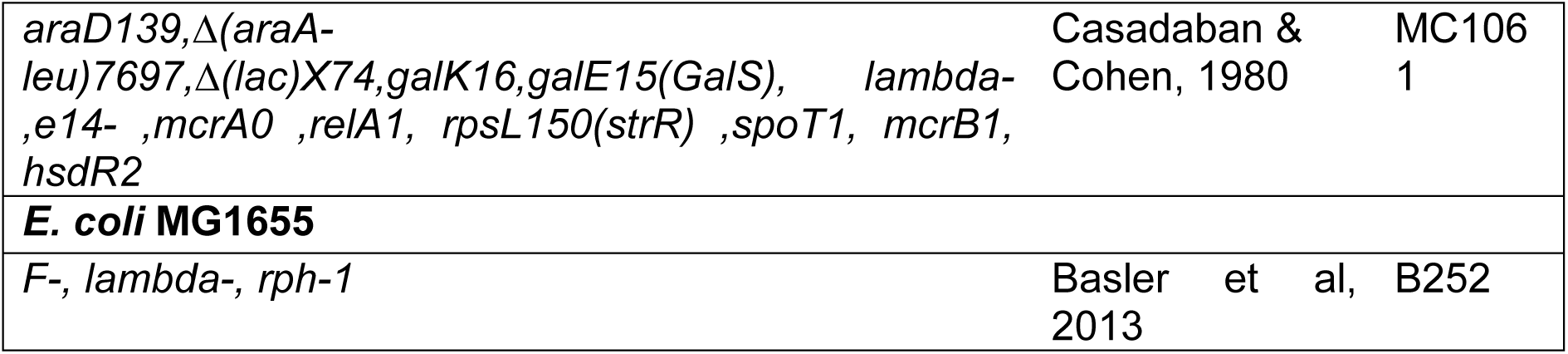
Strains used in this study.

**Supplementary Table 2:**
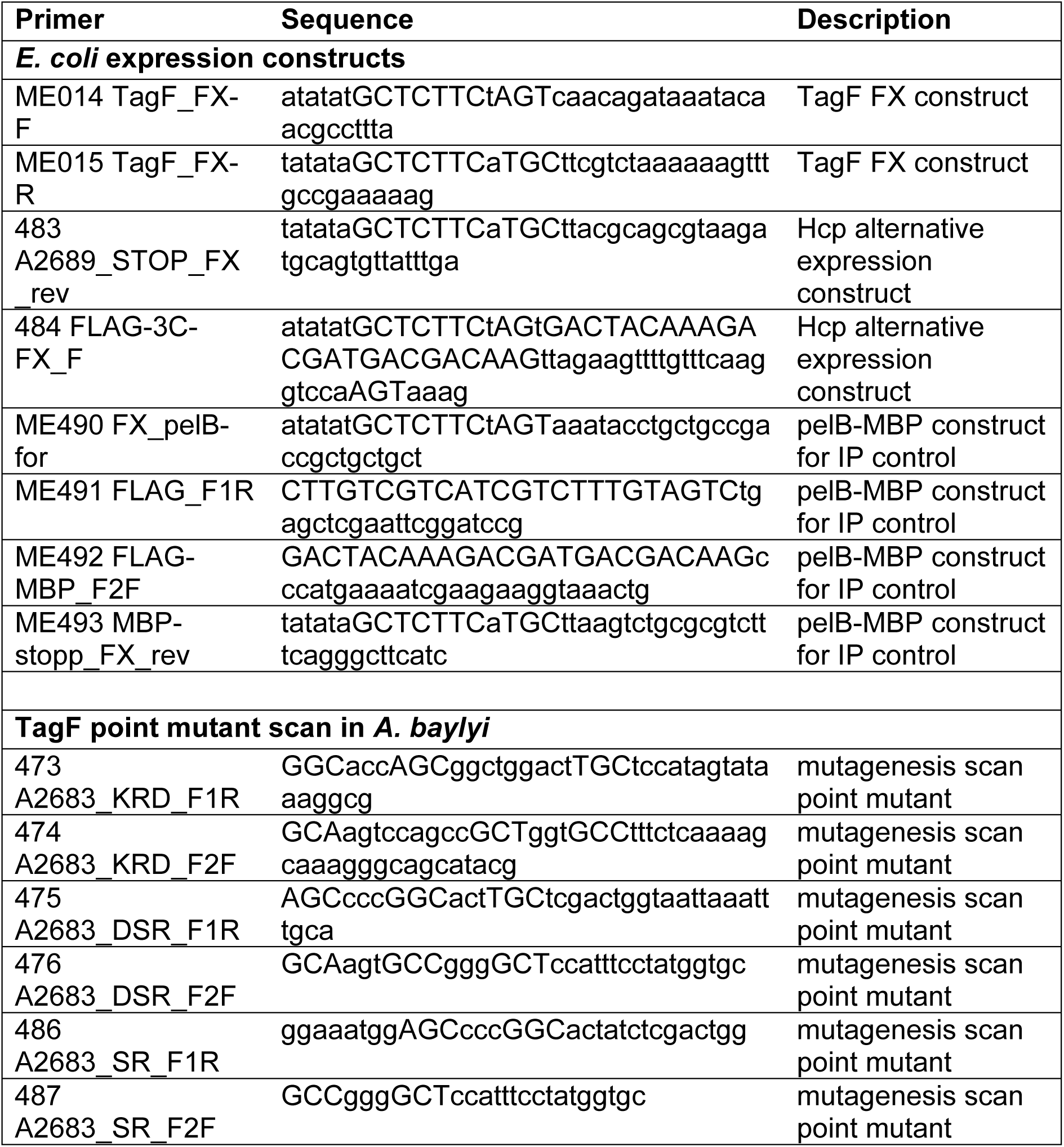
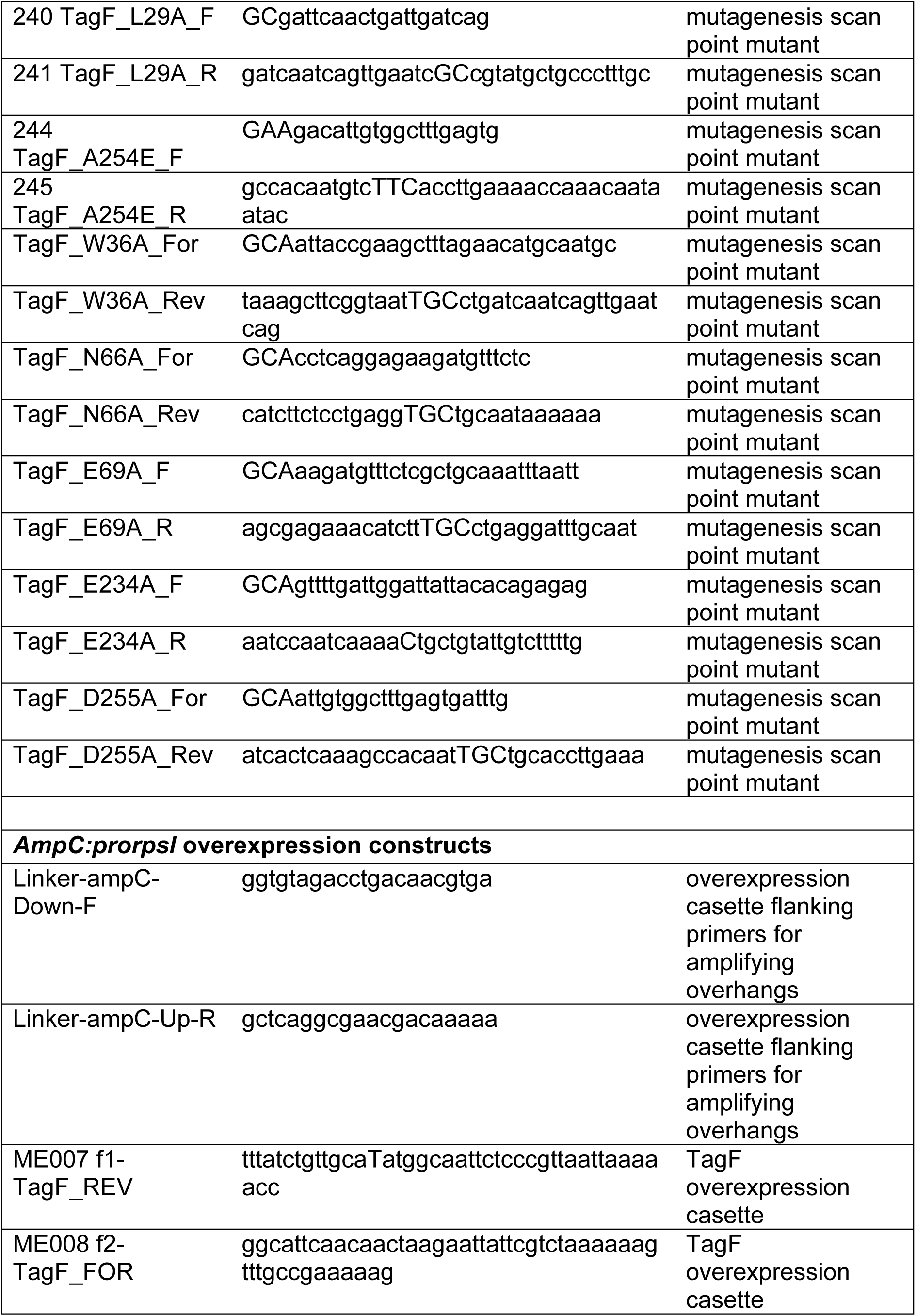
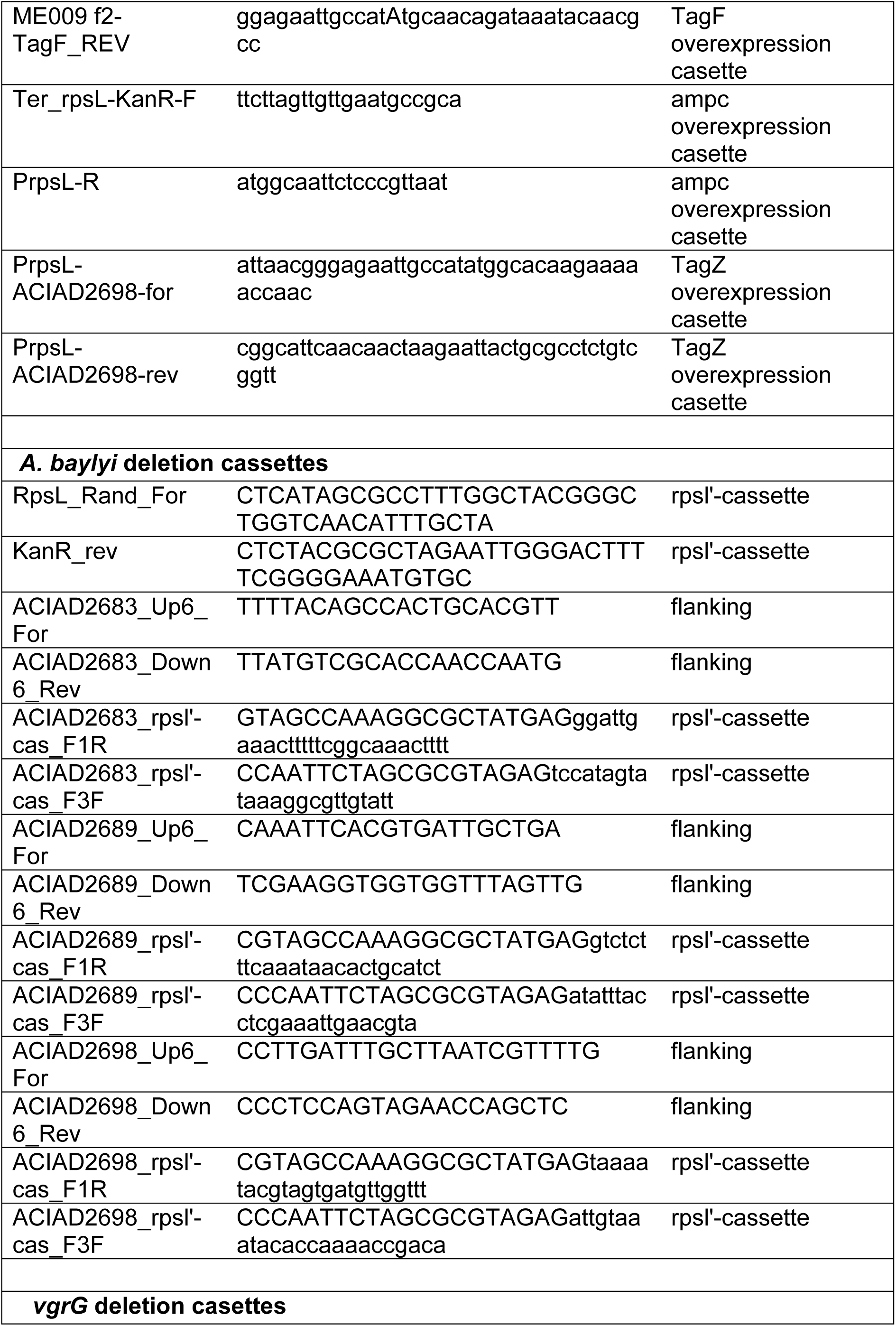
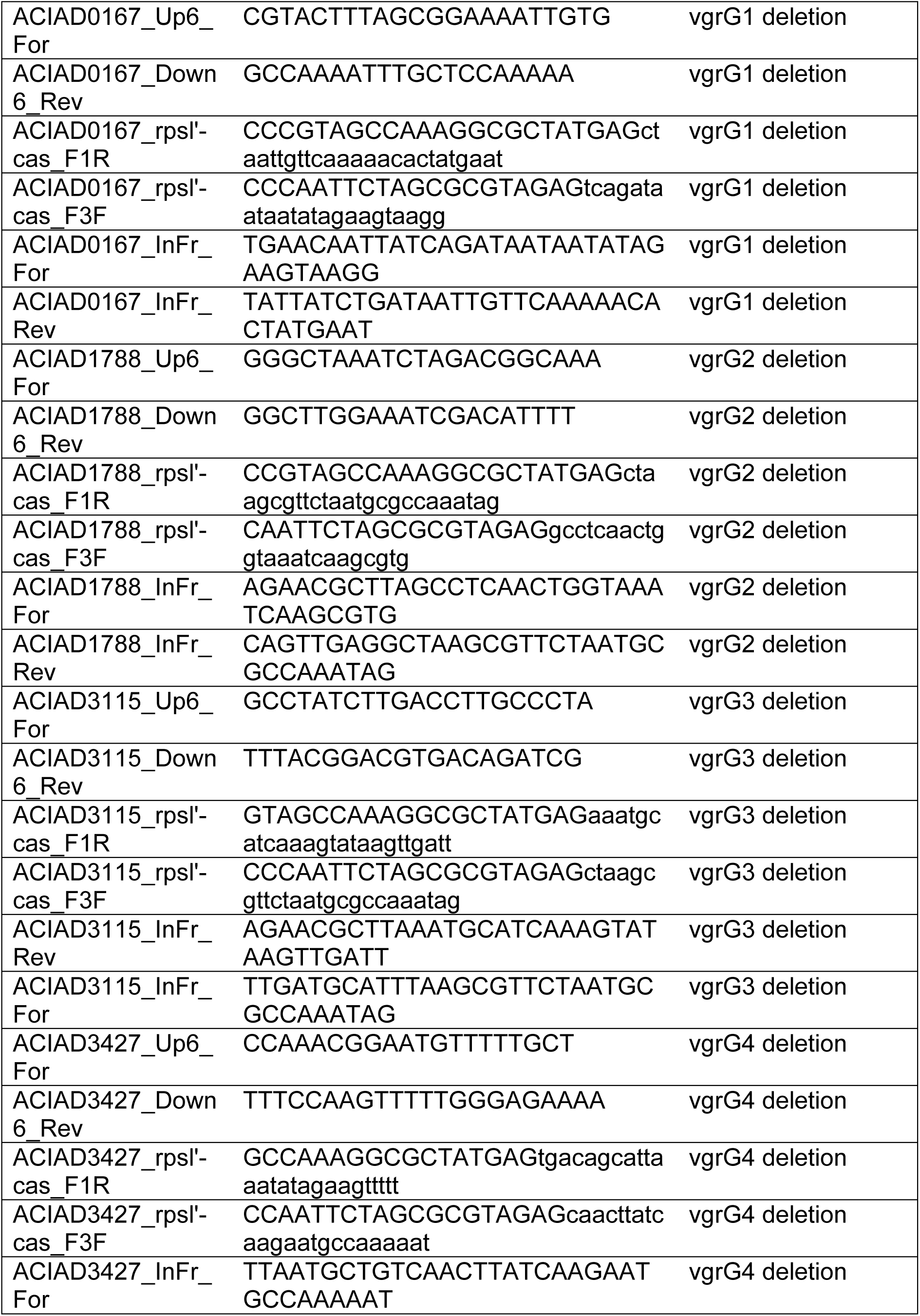
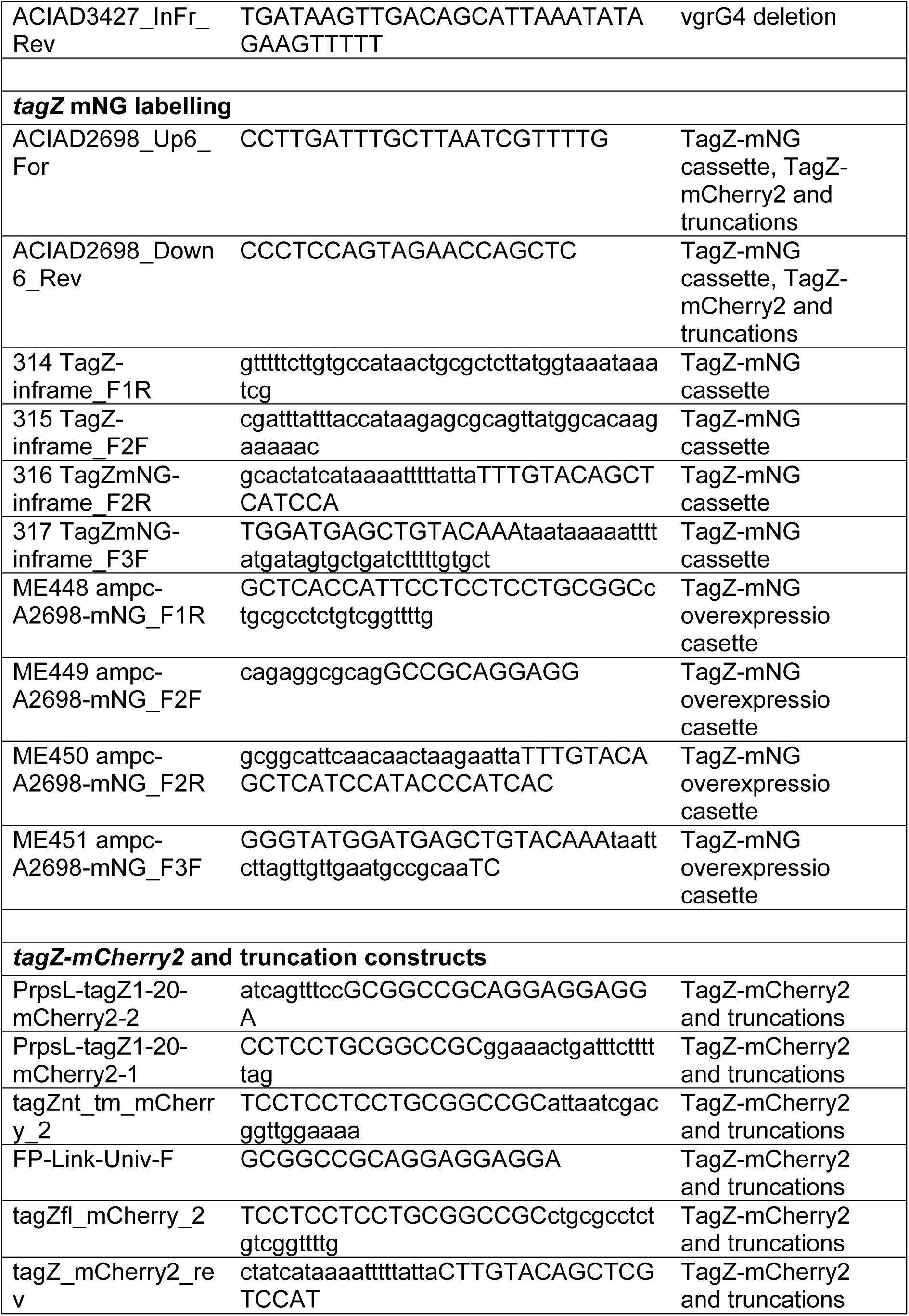
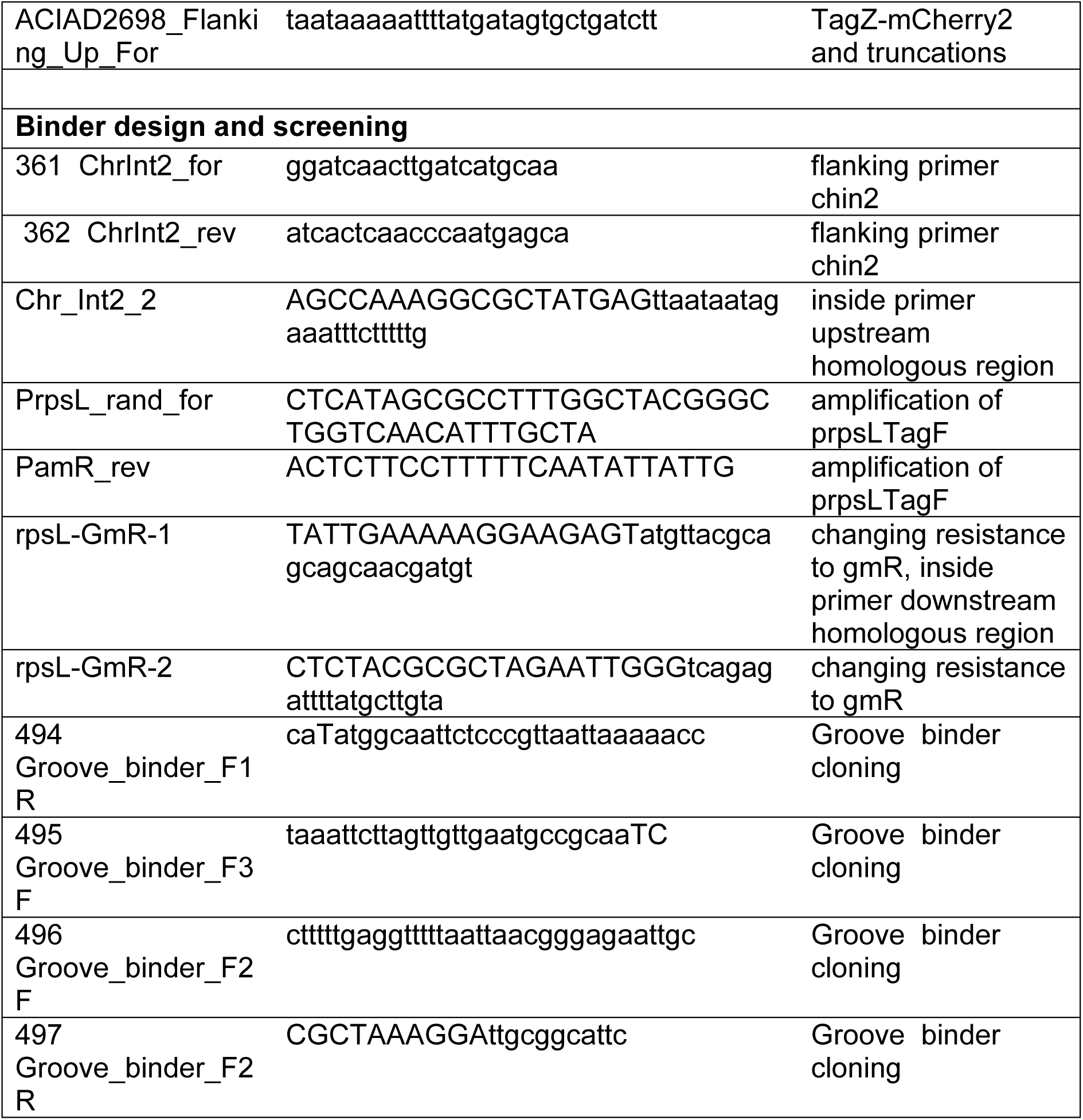
Oligonucleotides used in this study.

**Supplementary Table 3:** Plasmids used in this study.

| Genotype | Resistance | Reference | Plasmid ID |
| --- | --- | --- | --- |
| pBXC3H-*tagF* | Ampicilin,<br>150 ug/ml | This study | ME028 |
| pBXC3H_FLAG-3C-ACIAD2689_stop |  |  | ME469 |
| pBX-pelB-His<sub>10</sub>-mbp |  |  | ME490 |
| pBX-pelB-FLAG-mbp |  |  | ME491 |
| pBXC3H-*tagF*(K13A+R17AD19A) |  |  | ME473 |
| pBXC3H-*tagF*(K13A+R17AD19A<br>+S84A+R86A) |  |  | ME499 |
| pBXC3H-*tagF*(E234A) |  |  | ME477 |
| pInit.cat | Chloramphenico<br>l, 35 µg/ml | (Geertsma,<br>2013) | ME010 |
| pBXC3H | Ampicilin,<br>150 ug/ml | (Geertsma,<br>2013) | ME012 |
| pBXNPH3-M3 | Ampicilin,<br>150 ug/ml | Geertsma, 2013 | ME017 |

**Supplementary Table 4:** Antibodies used in this study.

| Antigen | Source | Supplier | Working dilution<br>in<br>5% milk PBST |
| --- | --- | --- | --- |
| Anti-TagF polyclonal | rabbit | Genscript | 1 ug/ml |
| anti-His<sub>6</sub>-HRP Monoclonal<br>A7058 | mouse | Sigma Aldrich | 1:20,000 |
| ANTI-FLAG® M2 Monoclonal<br>F1804 | mouse | Sigma-Aldrich | 1:1,000 |
| Anti-Strep Tag II Monoclonal<br>A5D10 | mouse | Thermo Fisher<br>Scientific | 10 µg/ml |
| Peroxidase AffiniPure Goat Anti-Mouse IgG (H+L) | goat | Jackson<br>Immuno<br>Research LTD | 1:5,000 |
| Peroxidase AffiniPure Goat Anti-Rabbit IgG (H+L) | goat | Jackson<br>Immuno<br>Research LTD | 1:5,000 |

## Supplementary Files

**Supplementary File 1:** TagF GCSnap output and files detailing relative frequency of gene occurrence in the data as used for Figure 1A.

**Supplementary File 2:** TagF phylogenetic analysis, including the fully labelled and annotated tree corresponding to Figure 1B, Supplementary Figure 2.

**Supplementary File 3:** TagF and TagZ alignment files regarding Figure 1B, Figure 3B, and Figure 4B.

**Supplementary File 4:** Proteomics results for TagF overexpression and co-purification experiments.

